# The key protein of endosomal mRNP transport Rrm4 binds translational landmark sites of cargo mRNAs

**DOI:** 10.1101/418079

**Authors:** Lilli Olgeiser, Carl Haag, Susan Boerner, Jernej Ule, Anke Busch, Janine Koepke, Julian König, Michael Feldbrügge, Kathi Zarnack

## Abstract

RNA-binding proteins (RBPs) determine spatiotemporal gene expression by mediating active transport and local translation of cargo mRNAs. Here, we cast a transcriptome-wide view on the transported mRNAs and cognate RBP binding sites during endosomal messenger ribonucleoprotein (mRNP) transport in *Ustilago maydis*. Using individual-nucleotide resolution UV crosslinking and immunoprecipitation (iCLIP), we compare the key transport RBP Rrm4 and the newly identified endosomal mRNP component Grp1 that is crucial to coordinate hyphal growth. Both RBPs bind predominantly in the 3’ untranslated region of thousands of shared cargo mRNAs, often in close proximity. Intriguingly, Rrm4 precisely binds at stop codons, which constitute landmark sites of translation, suggesting an intimate connection of mRNA transport and translation. Towards uncovering the code of recognition, we identify UAUG as specific binding motif of Rrm4 that is bound by its third RRM domain. Altogether, we provide first insights into the positional organisation of co-localising RBPs on individual cargo mRNAs.

## Introduction

All eukaryotic cells must accurately regulate the expression of proteins in time and space. To this end, many mRNAs accumulate at specific subcellular sites, and their local translation is exactly timed [1-3]. mRNA localisation is achieved most commonly by active motor-dependent transport along the cytoskeleton. Functional transport units are messenger ribonucleoprotein complexes (mRNPs), consisting of various RNA-binding proteins (RBPs), accessory proteins and cargo mRNAs. Key factors are RBPs that recognise localisation elements (LEs) within mRNAs. For movement, the RBPs either interact with motors directly or are connected via linker proteins [1, 4].

We discovered co-transport of mRNPs on the cytoplasmic surface of early endosomes as a novel translocation mechanism of cargo mRNAs during hyphal growth in fungi [5, 6]. These endosomes shuttle along microtubules by the concerted action of plus-end directed kinesin and minus-end directed dynein [7, 8]. They serve as multipurpose platforms functioning not only during endocytic recycling but also during long-distance transport of whole organelles such as peroxisomes [9-12].

Endosomal mRNA transport was uncovered analysing the RBP Rrm4 in the dimorphic phytopathogenic fungus *Ustilago maydis* (Fig EV1A) [13, 14]. Loss of Rrm4 has no effects on the yeast form of the fungus. However, the absence of Rrm4 causes characteristic defects in unipolar growth when switching to the hyphal form: the fraction of bipolarly growing hyphae increases and the insertion of basal septa is delayed [6, 15]. In line with endosomal mRNA transport, Rrm4 binds mRNAs and shuttles on early endosomes along microtubules *in vivo* [5, 16]. Using the poly(A)-binding protein Pab1 as an mRNA marker revealed that loss of Rrm4 abolishes this transport, resulting in a gradient of mRNAs declining from the nucleus towards the cell periphery [17]. Thus, one function of Rrm4 might be the general distribution of mRNAs within hyphae [17].

Initial CLIP experiments with Rrm4 identified target mRNAs encoding chitinase Cts1 and septin Cdc3, among others [17, 18]. The subcellular localisation of both proteins was Rrm4-dependent: Loss of Rrm4 strongly reduced the secretion of the chitinase Cts1. Moreover, shuttling of the Cdc3 protein on early endosomes was abolished, and the gradient of septin filaments at the growth pole of hyphae was no longer formed [6]. Since *cdc3* mRNA and its encoded protein are found together with ribosomes on the same shuttling endosomes, we hypothesised that endosome-coupled translation of *cdc3* mRNA during long-distance transport is critical for the efficient formation of septin filaments at the growth pole [6]. This was supported by demonstrating that all four septin-encoding mRNAs are present on endosomes and that septin proteins assemble into heteromeric complexes on the cytoplasmic face of endosomes during long-distance transport [19]. Thus, Rrm4-dependent mRNA transport regulates the specific localisation of the corresponding translation products. To understand this complex process at the transcriptome-wide level, we present herein an *in vivo* snapshot of RNA binding sites of endosomal RBPs on cargo mRNAs at single-nucleotide resolution.

## Results

### Loss of the glycine/glutamine-rich protein Grp1 affects hyphal growth

In order to identify additional protein components involved in endosomal mRNA transport, we performed pilot affinity tag purification using Rrm4 as bait. We identified the potential RBP glycine-rich protein 1 (Grp1; UMAG_02412), which carries an N-terminal RNA recognition motif (RRM) domain followed by a short C-terminal region rich in glycine and glutamine (GQ-rich; Fig 1A, Fig EV1B-D). The protein was similar to other small RRM proteins, such as human CIRBP or RBM3 and plant RBG7 (*At*GRP7), all previously described as global stress regulators (Fig 1A) [20, 21].

**Fig 1.**
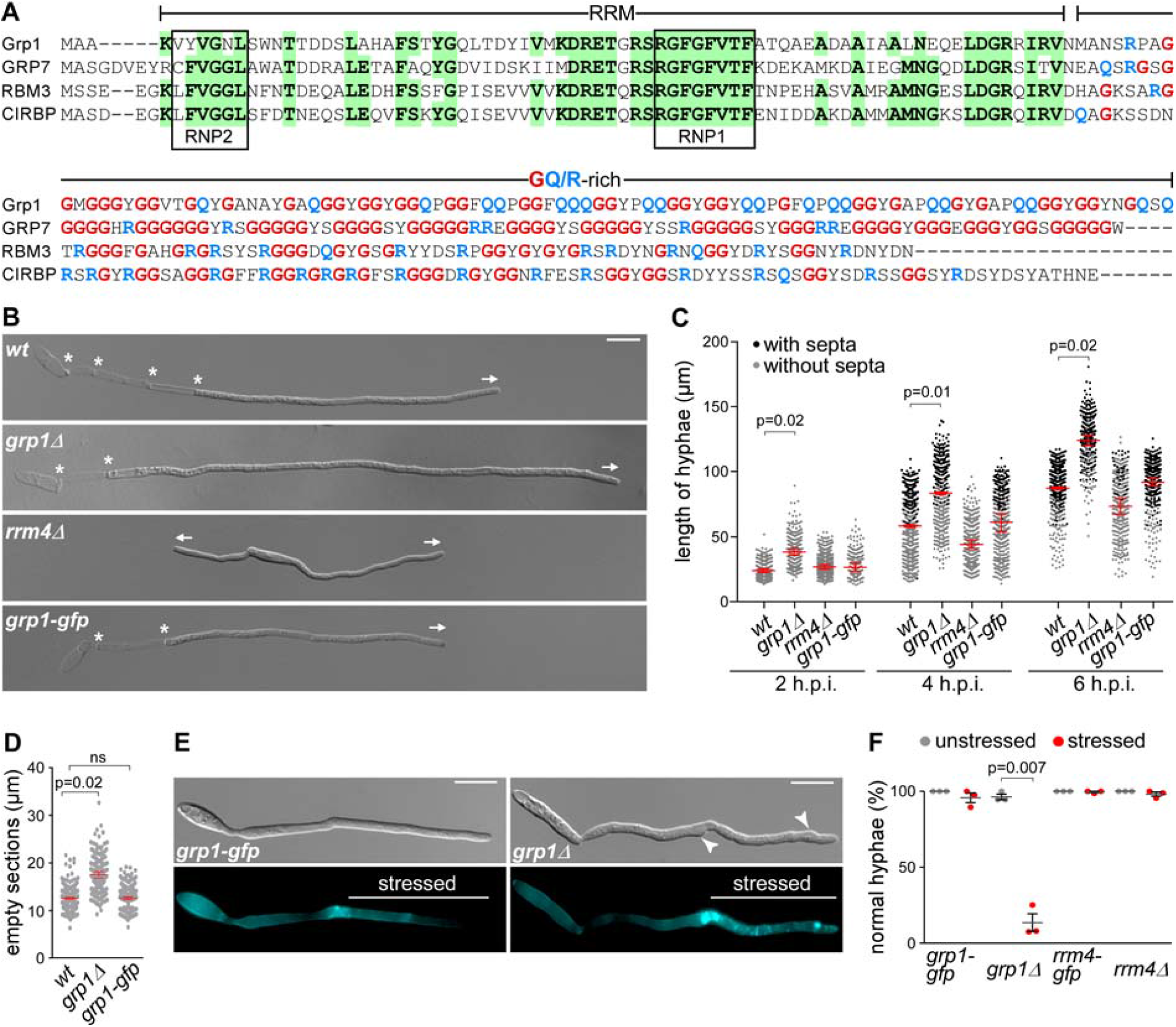
Grp1 is important for hyphal growth under suboptimal conditions. (**A**) Sequence alignment of glycine-rich proteins (Fig EV1B). *Um*Grp1 from *U. maydis* (UMAG_02412), *At*GRP7 (RBG7) from *A. thaliana* (NC_003071.7), *Hs*RBM3 and *Hs*CIRBP from *H. sapiens* (NC_000023.11 and NC_000019.10, respectively). Amino acid positions within RRM that are identical in at least three proteins are highlighted in green (boxes indicate RNA contact regions RNP1 and RNP2). Glycine and arginine/glutamine residues in the glycine-rich region are labelled in red and blue, respectively. (**B**) Hyphae of AB33 derivatives (6 h.p.i.). Growth direction and basal septa are marked by arrows and asterisks, respectively (scale bar, 10 μm). (**C**) Hyphal length over time. Black and grey dots represent hyphae with and without septa, respectively. Shown are merged data (> 200 hyphae per strain) from three independent experiments, overlaid with the mean of means, red line, and standard error of the mean (s.e.m.). Significance was assessed using paired two-tailed Student’s t-test on the mean hyphal lengths from the replicate experiments, followed by multiple testing correction (Benjamini-Hochberg). Significant p-values (p < 0.05) for comparison against wild type within each time point are indicated above. (**D**) Length of empty sections (see Fig EV1J). Merged data from three independent experiments are shown together, overlaid with mean of means, red line, and s.e.m. (total hyphae analysed: *wt*, 250; *grp1Δ*, 190; *grp1-gfp*, 332; difference in means between *wt* and *grp1-gfp* was statistically not significant, ns, p= 0.95; paired two-tailed Student’s t-test on the mean lengths from the replicate experiments (n=3). (**E**) Differential interference contrast (DIC, top) and fluorescence images (bottom) of AB33 hyphae (5 h.p.i.) stressed at 3 h.p.i. with cell wall inhibitor CFW (2.5 μM). Arrowheads indicate aberrant cell wall deformation (scale bar, 10 μm). (**F**) Percentage of hyphae with normal cell walls with (stressed) and without (unstressed) CFW treatment (data points represent percentages of three independent experiments, n=3; mean, dark grey lines, and standard error of the mean, s.e.m., > 100 hyphae analysed per experiment; paired two-tailed Student’s t-test on the means).

For functional analysis, we generated deletion mutants in laboratory strain AB33. In this strain the master transcription factor controlling hyphal growth is under control of an inducible promoter. Thus, hyphal growth can be elicited synchronously by changing the nitrogen source in the medium. The corresponding hyphae grow like wild type by tip expansion at the apical pole, while the nucleus is positioned in the centre and septa are inserted in regular intervals at the basal pole (Fig EV 1A) [22]. In the yeast form of AB33, we observed that loss of Grp1 resulted in slower proliferation as well as increased cell size (Fig EV1E-G). At lower temperatures, growth of the *grp1Δ* strain was affected even more strongly and it exhibited an altered colony morphology (Fig EV1H). This was consistent with a potential function in cold stress response, similar to the plant and human orthologues [20, 21]. Furthermore, colony growth of the *grp1Δ* strain was strongly reduced upon treatment with inhibitors of cell wall biosynthesis, such as Calcofluor White (CFW) or Congo Red (CR) [23, 24]. Hence, loss of Grp1 might cause defects in cell wall formation (Fig EV1I).

Studying hyphal growth revealed that, unlike observed in *rrm4Δ* strains, loss of Grp1 did not cause an increased amount of bipolar cells as it is characteristic for defects in microtubule-dependent transport (see *rrm4Δ* hyphae for comparison; Fig 1B-C) [12]. On the contrary, under optimal growth conditions hyphae were significantly longer (Fig 1B-C), and the length of empty sections at the basal pole was increased (Fig 1D; Fig EV1J). Hence, the coordination of hyphal growth may be disturbed in the absence of Grp1. In order to further investigate this, we stressed hyphae three hours post induction (h.p.i.) of hyphal growth with the cell wall inhibitor CFW. In comparison to wild type hyphae, we observed a strongly increased number of *grp1Δ* hyphae with abnormal shapes (86%), indicating that cell wall integrity might be affected (Fig 1E-F).

In summary, loss of Grp1 affects both yeast-like and hyphal growth. During the latter, Grp1 seems to be crucial for the correct coordination of cell wall expansion, which becomes particularly apparent during stress conditions.

### Grp1 is a novel component of endosomal mRNA transport

To analyse the subcellular localisation of Grp1, we generated AB33 strains expressing Grp1 fused at its C-terminus to Gfp by homologous recombination. The functional Grp1-Gfp version accumulated in the cytoplasm as well as in the nucleus of hyphae. In comparison, the poly(A)-binding protein Pab1-Gfp was absent from the nucleus, suggesting that this localisation pattern is specific for Grp1 (Fig 2A). Importantly, a subpopulation of Grp1-Gfp moved bi-directionally in the cytoplasm with a velocity comparable to Rrm4-Gfp and Pab1-Gfp, which are known to shuttle on early endosomes (Fig 2A-B; Supplemental Video 1).

**Fig 2.**
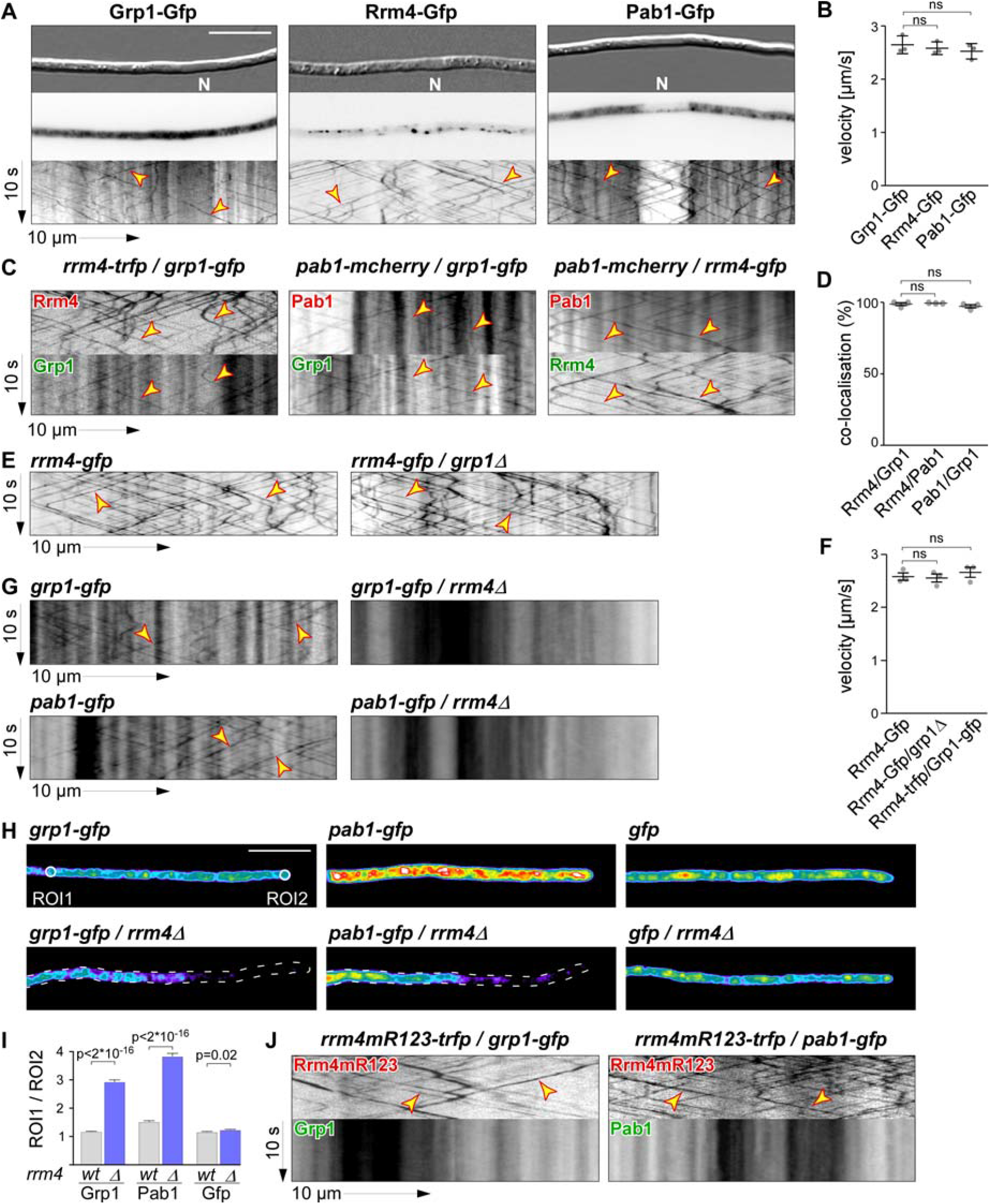
Grp1 shuttles on Rrm4-positive endosomes throughout hyphae. (**A**) Micrographs (DIC and inverted fluorescence image; scale bar, 10 μm) and corresponding kymographs of AB33 hyphae (6 h.p.i.) expressing Grp1-Gfp, Rrm4-Gfp or Pab1-Gfp (arrow length on the left and bottom indicate time and distance, 10 s and 10 μm, respectively). To visualise directed movement of signals (distance over time) within a series of images, kymographs were generated by plotting the position of signals along a defined path (x-axis) for each frame of the corresponding video (y-axis). Bidirectional movement is visible as diagonal lines (yellow arrowheads; N, nucleus; Supplemental Video 1). For an example image of a complete AB33 hypha, see Fig EV1A. (**B**) Average velocity of fluorescent signals per hypha for strains shown in A (movement of tracks with > 5 μm were scored as processive). Data points represent averages from three independent experiments (n=3), with mean, red line, and s.e.m‥ At least 20 signals/hyphae were analysed out of 12 hyphae per experiment (ns; p=0.18 and p=0.23) using an paired twotailed Student’s t-test on the means. (**C**) Kymographs of hyphae of AB33 derivatives (6 h.p.i.) expressing pairs of red and green fluorescent proteins as indicated. Fluorescence signals were detected simultaneously using dual-view technology (arrow length as in A). Processive co-localising signals are marked by yellow arrowheads (Supplemental Videos 2-4). (**D**) Percentage of red fluorescent signals exhibiting co-localisation with the green fluorescent signal for strains shown in (C). Data points represent observed co-localisation in three independent experiments, mean, dark grey line, and s.e.m. (n=3, 11 hyphae each; paired twotailed Student’s t-test on the means; ns; p=0.63 and p=0.5). (**E**) Kymographs comparing hyphae of AB33 derivatives (6 h.p.i.) expressing Rrm4-Gfp in the wild type (left) or *grp1Δ* strains (right) (processive signals marked by yellow arrowheads; arrow length on the left and bottom indicate time and distance, 10 s and 10 μm, respectively; Supplemental Video 5). (**F**) Average velocity of fluorescent signals per hyphae for strains shown in (E) (movement of tracks with > 5 μm were scored as processive). Data points represent averages from three independent experiments (n=3) with mean, black line, and s.e.m‥ At least 20 signals/hypha were analysed out of at least 10 hyphae per experiment (ns; p=0.27 and p=0.4) using an paired two-tailed Student’s t-test on the means. (**G**) Kymographs comparing hyphae (6 h.p.i.) expressing Grp1-Gfp or Pab1-Gfp in wild type (left) with *rrm4Δ* strains (right; processive signals marked by yellow arrowheads; arrow length as in A). (**H**) Hyphal tips (4 h.p.i.) of AB33 derivatives expressing Grp1-Gfp, Pab1-Gfp or Gfp alone comparing wild type (top) with *rrm4Δ* strains (bottom). Fluorescence micrographs in false colours (black/blue to red/white, low to high intensities, respectively; scale bar, 10 μm; ROI1 and ROI2-labelled circles exemplarily indicate regions-of-interest analysed in E). (**I**) Ratio of signal intensities in strains shown in (H) comparing Gfp fluorescence at the tip (ROI1) and in close vicinity to the nucleus (ROI2; see Materials and methods). Bars represent mean and s.e.m. (*wt*: n=160; *rrm4Δ*: n=152), Pab1G (n=126; *rrm4Δ*: n=170), Gfp (n=152; *rrm4Δ*: n=210), unpaired two-tailed Student’s t-test. (**J**) Kymographs of hyphae of AB33 derivatives (6 h.p.i.) expressing pairs of red and green fluorescent proteins as indicated (arrow length as in A; Supplemental Videos 6-7). Fluorescence signals were detected simultaneously using dual-view technology. Processive co-localising signals are marked by yellow arrowheads. Note that processive movement is completely lost in the lower panels. Only static signals, visualised as vertical lines, are remaining.

To test whether Grp1 shuttles on Rrm4-positive endosomes, we performed dynamic co-localisation studies using dual-view technology [25]. We generated AB33 strains co-expressing Grp1-Gfp and Rrm4 fused C-terminally to the red fluorescent protein Tag-Rfp (tRfp) [26]. For comparison, we used a strain expressing Pab1 fused to the red fluorescent protein mCherry [17, 27]. Analysing hyphae 6 h.p.i. 99% of processive Grp1-Gfp signals co-migrated with Rrm4-tRfp, revealing extensive co-localisation of both proteins in shuttling units (Fig 2C-D; Supplemental Video 2). Consistently, 97% of processive Grp1-Gfp signals co-migrated with Pab1-mCherry, indicating that Grp1, like Pab1, was present on Rrm4-positive endosomes (Fig 2C-D; Supplemental Videos 3-4). Thus, Grp1 appears to be a novel component of endosomal mRNPs that might already be recruited to transport mRNPs in the nucleus.

### The endosomal localisation of Grp1 depends on Rrm4

To investigate whether Grp1 has an influence on the shuttling of Rrm4-positive endosomes, we studied Rrm4 movement in *grp1Δ* strains. Loss of Grp1 altered neither processive Rrm4-Gfp movement nor the velocity of the respective endosomes (Fig 2E-F; Supplemental Video 5). Vice versa, studying Grp1-Gfp movement in the absence of Rrm4 revealed that its endosomal localisation depended on Rrm4 (Fig 2G). Importantly, similar to Pab1-Gfp, a gradient of Grp1-Gfp was formed in *rrm4Δ* hyphae, with a decreasing signal intensity towards the growing apex (Fig 2H-I) [17]. Similar to Pab1, which is expected to associate with almost all poly(A) tails of mRNAs [28], Grp1 might therefore be distributed in association with many mRNAs (see below).

To test whether Grp1-Gfp binds to endosomes in an mRNA-dependent manner, we generated AB33 strains expressing Rrm4^mR123^-tRfp. This Rrm4 variant carried point mutations in the RNP1 regions of RRM domains 1-3 causing a reduced RNA binding activity and loss of function of Rrm4 [16]. In dual-view experiments we observed that Grp1-Gfp like Pab1-Gfp no longer shuttled in the presence of Rrm4^mR123^-tRfp (Fig 2J). Thus, the localisation of Grp1 depends on the presence of functional Rrm4, more precisely on its capability to bind RNA. In summary, we identified Grp1 as a novel component of endosomal mRNPs whose shuttling on Rrm4-positive endosomes depended on Rrm4 and mRNA.

### Rrm4 and Grp1 share thousands of target transcripts

In order to learn more about the function of the two endosomal mRNP components Rrm4 and Grp1 during hyphal growth, we performed a comparative transcriptome-wide analysis of their RNA binding behaviour using individual-nucleotide resolution UV crosslinking and immunoprecipitation (iCLIP) [29]. For application with fungal RBPs, we had to modify a number of steps in the iCLIP protocol (Fig EV2A-D; see Materials and methods) [30]. One major challenge was the high RNase and protease activity in fungal cell extracts that resulted in a low yield of crosslinked protein-RNA complexes and short mRNA fragments. The most critical changes to the protocol came with the fast processing of crosslinked material and the identification of the optimal UV-C irradiation dose (Fig EV2B).

Using the improved protocol, we found that Rrm4-Gfp and Grp1-Gfp displayed substantial crosslinking to RNA *in vivo* (compared to Gfp control; Fig 3A). As expected, the RNA signal was dependent on UV-C irradiation and sensitive to RNase I digestion. Upon iCLIP library preparation, we obtained more than 100 million sequencing reads, corresponding to 4.7 x 10^6^ and 14.8 x 10^6^ crosslink events for Rrm4 and Grp1, respectively (Fig EV3A-B). Reproducibility between two replicate experiments was high for both proteins, demonstrating the quality of the obtained data set (Pearson correlation coefficient > 0.96, p value < 2.22e-16; Fig EV3C).

**Fig 3.**
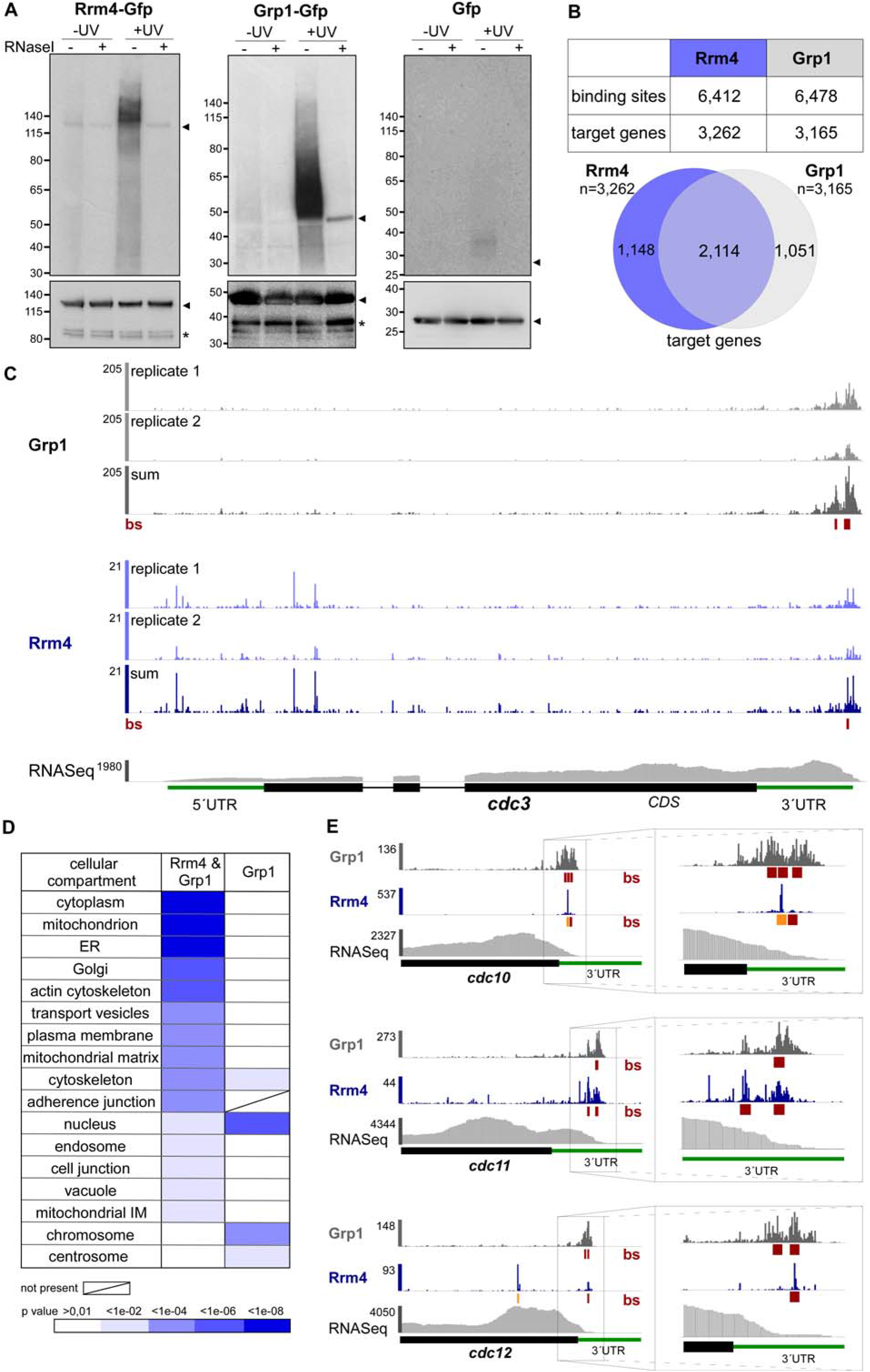
Rrm4 and Grp1 bind to thousands of target transcripts. (**A**) Autoradiograph and Western blot analyses for representative iCLIP experiments with Rrm4-Gfp, Grp1-Gfp and Gfp. Upper part: Upon radioactive labelling of co-purified RNAs, the protein-RNA complexes were size-separated in a denaturing polyacrylamide gel. Protein-RNA complexes are visible as smear above the size of the protein (Rrm4-Gfp, 112 kDa; Grp1-Gfp, 45 kDa; Gfp, 27 kDa; indicated by arrowheads on the right). Samples with and without UV-C irradiation and RNase I (see Material and methods) are shown. Lower part: corresponding Western blot analysis using α-Gfp antibody (arrowheads and asterisks indicate expected protein sizes and putative degradation products, respectively). (**B**) Summary of binding sites and target transcripts of Rrm4 and Grp1 (top). Venn diagram (below) illustrates the overlap of Rrm4 and Grp1 target transcripts. (**C**) iCLIP data for Rrm4 and Grp1 on *cdc3* (UMAG_10503; crosslink events per nucleotide from two experimental replicates [light grey/light blue] and merged data [grey/blue] from AB33 filaments, 6 h.p.i.). Track below the merged iCLIP data show binding sites for each protein (bs, red). Note that crosslink events in the 5’ UTR and first exon of *cdc3* were not assigned as binding sites due to low reproducibility between replicates (see Materials and methods). RNASeq coverage from wild type AB33 filaments (6 h.p.i.) is shown for comparison. Gene model with exon/intron structure below was extended by 300 nt on either side to account for 5’ and 3’ UTRs (green). (**D**) Functional categories of cellular components (FunCat annotation scheme, 78) for proteins encoded by target transcripts that are shared between Rrm4 and Grp1 (left) or unique to Grp1 (right). P values for the enrichment of the listed category terms are depicted by colour (see scale below). (**E**) iCLIP data (crosslink events from merged replicates) for Rrm4 (blue) and Grp1 (grey) as well as RNASeq coverage on selected target transcripts (*cdc10*, UMAG_10644; *cdc11*, UMAG_03449; *cdc12*, UMAG_03599). Enlarged regions (indicated by boxes) of the 3’ UTR (green) are shown on the right. Datasets and visualisation as in (C). Binding sites (bs) are shown (red; orange indicates overlap with UAUG).

Consistent with the abundance of both proteins, the crosslink events accumulated into thousands of clusters that spread across major parts of the transcriptome (Fig EV3A). In order to focus on the most prominent sites, we used the crosslink frequency within each cluster relative to the background signal within the same transcript to determine the 25% most prominent binding sites for Rrm4 and Grp1 (‘signal-over-background’; see Materials and methods). This procedure identified a total of 6,412 binding sites for Rrm4 and 6,478 binding sites for Grp1, residing in 3,262 and 3,165 target transcripts, respectively (Fig 3B). This represented a substantial fraction of the about 6,700 protein-encoding genes in the *U. maydis* genome [31]. Extensive endosomal transport of mRNA is consistent with a role in evenly distributing mRNAs throughout hyphae (see Discussion).

Comparing Rrm4 and Grp1 revealed a large overlap of 2,114 target transcripts that were conjointly bound by both proteins (Fig 3B-C, Dataset EV1; see below). In this shared target set, we observed an enrichment for functional categories like mitochondrion, vesicle transport and cytoskeleton (Fig 3D). Moreover, we found several known Rrm4 target transcripts, including for instance all four *septin* mRNAs (Fig 3C, 3E). Binding sites of Rrm4 and Grp1 in the *septin* mRNAs were almost exclusively located in the 3’ untranslated region (UTR), consistent with the hypothesis that these mRNAs are transported in a translationally active state (see Discussion) [19]. *cts1* mRNA, another known target of the Rrm4 transport machinery [18], also carried binding sites of both RBPs in the 3’ UTR (Fig EV3D).

To assess the function of target mRNAs that are specifically recognised by only one of the two RBPs, we applied more stringent criteria to define 280 and 520 transcripts that were uniquely bound by Rrm4 and Grp1, respectively (Fig EV3E, Dataset EV2-3). While the Rrm4-unique set displayed no clear trend, the Grp1-unique set showed an enrichment for mRNAs encoding nuclear proteins that were involved in transcriptional regulation and chromatin remodelling (Fig 3D). Although these mRNAs were expressed and bound by Grp1, they were most likely not transported by the Rrm4 machinery, which we presume would result in perinuclear localisation. This could facilitate an efficient nuclear import of the translation products, as being described in mammalian cells for transcription factors like c-Myc or for metallothionin [32, 33].

In summary, the comparative iCLIP approach revealed that Grp1 and Rrm4 conjointly bind thousands of shared target mRNAs. These offer a comprehensive view on the full spectrum of cargo mRNAs transported by the endosomal mRNP transport machinery in *U. maydis*.

### Rrm4 binds to functionally important sites of target transcripts

Studying the distribution of binding sites in different transcript regions revealed that Rrm4 and Grp1 preferentially bound in the 3’ UTR (Fig 4A). Within this region, both proteins frequently bound in close proximity, with 51% of Rrm4 binding sites directly overlapping with a Grp1 binding site (compared to only 5% in the open reading frame, ORF; Fig 4B-C). Thus, the cargo mRNAs of the transport mRNPs are often conjointly recognised by both RBPs in the 3’ UTR.

**Fig 4.**
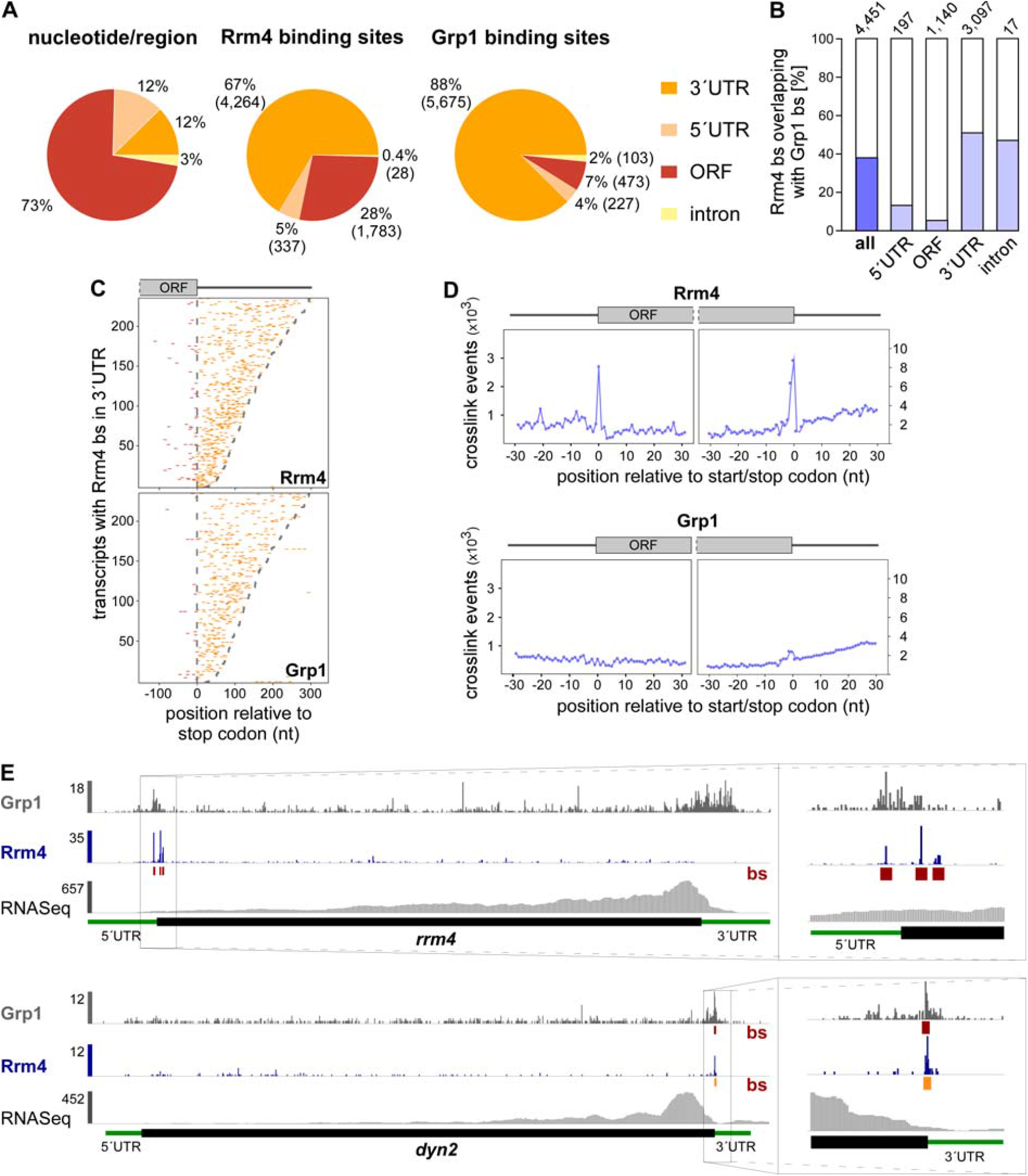
Rrm4 binds target transcripts at the start and stop codon. (**A**) Distribution of binding sites within different transcript regions: 5’ UTR, 3’ UTR, ORF and intron. Percentage and absolute number of binding sites are given for each category. On the left, a transcriptome-wide distribution of nucleotides per transcript region is shown for comparison. (**B**) Percentage of Rrm4 binding sites (bs) overlapping with Grp1 bs (by at least 1 nt) within shared target transcripts, shown for all bs and separated into transcript regions. The total number of binding sites per category is indicated on top. (**C**) Positional maps of Rrm4 (top) and Grp1 (bottom) bs relative to the stop codon (position 0). Binding sites in ORFs and 3’ UTR are given in red and orange, respectively. 234 target transcripts were randomly selected carrying an Rrm4 bs in the 3’ UTR (with > 100 Rrm4 crosslink events; out of 1,715 Rrm4/Grp1 shared targets with Rrm4 bs in 3’ UTR; Dataset EV6). Transcripts were ordered by decreasing 3’ UTR length (see Materials and methods). (**D**) Metaprofiles of Rrm4 (top) and Grp1 (bottom) crosslink events relative to the start and stop codon (position 0). Note that crosslink events are substantially more frequent towards ORF ends, reflected in different y-axis scales. (**E**) Genome browser views of Rrm4 and Grp1 iCLIP events as well as RNASeq data of *sui1* (UMAG _02665) and *dyn2* (UMAG_04372). Visualisation as in Fig 3C.

In contrast to Grp1 that was almost exclusively attached to the 3’ UTR, Rrm4 bound a substantial fraction of target mRNAs within the ORF (1,315 mRNAs with 1,783 ORF binding sites; Fig 4A, 4C). Taking a closer look at the binding pattern of Rrm4 along ORFs, we observed binding sites at the start and stop codons of a subset of target mRNAs, reflected in increased crosslinking of Rrm4 at these sites (Fig 4D). Whereas only two transcripts harboured a Grp1 binding site at the start codon, Rrm4 binding sites overlapped the start codon in 47 target mRNAs, like in the transcript encoding the translation initiation factor Sui1 (UMAG_02665; Fig EV3D; Dataset EV4). Of note, the *rrm4* mRNA itself exhibited Rrm4 binding sites around the start codon, hinting at a potential autoregulation (Fig 4E).

Even more prominently than at start codons, we observed a strong accumulation of Rrm4 binding sites at the stop codons of multiple target transcripts (291 cases; Fig 4D-E; Dataset EV5). These included, for example, both subunits of cytoplasmic dynein (Dyn1 and Dyn2) [34]. Furthermore, the stop codon-bound targets were significantly enriched for mRNAs encoding mitochondrial proteins, including for instance the majority of nucleus-encoded subunits and accessory components of the F_O_F_1_-ATPase (Fig EV4A-C). Through binding at the stop codon, Rrm4 might influence translation termination for these targets. This hypothesis was supported by revisiting data from a previous proteomics approach analysing the membrane-associated proteome of wild type and *rrm4Δ* hyphae, in which the amount of the membrane-associated component Atp4 was threefold reduced in *rrm4Δ* hyphae compared to wild type (Fig EV4D) [18].

In essence, the high-resolution mapping of binding sites for the two endosomal RBPs Rrm4 and Grp1 revealed (i) that both proteins conjointly bind in the 3’ UTR, and (ii) that Rrm4 additionally recognises binding sites in the ORF and at start and stop codons.

### Rrm4 specifically recognises the motif UAUG via its third RRM

In order to address the RNA sequence specificity of both RBPs, we used the motif analysis algorithm DREME [35] to search for sequence motifs in a 30-nt window around the RBP binding sites. This analysis retrieved UAUG as the most significantly enriched sequence motif at Rrm4 binding sites (Fig 5A). Analysing the relative positioning of the motif showed that more than one third of all Rrm4 binding sites harboured a UAUG motif precisely at the centre of the binding site (2,201 out of 6,412; Fig 5B). The motif did not accumulate at Grp1 binding sites, supporting the notion that the motif was specifically recognised by Rrm4. In order to estimate the relative strength of Rrm4 binding, we calculated the ‘signal-over-background’ (SOB), i. e the ratio of crosslink events within the binding site over the background of crosslink events in the surrounding sequence (see Materials and methods). The background binding served as a proxy for the abundance of underlying transcript. Since the SOB procedure did not correct for UV crosslinking biases and similar confounding factors, comparisons between binding sites were only performed at a global scale. We observed that the Rrm4 binding sites with UAUG showed stronger relative binding than those lacking the motif (Fig 5C), suggesting a tight interaction of Rrm4 with the UAUG-associated binding sites.

**Fig 5.**
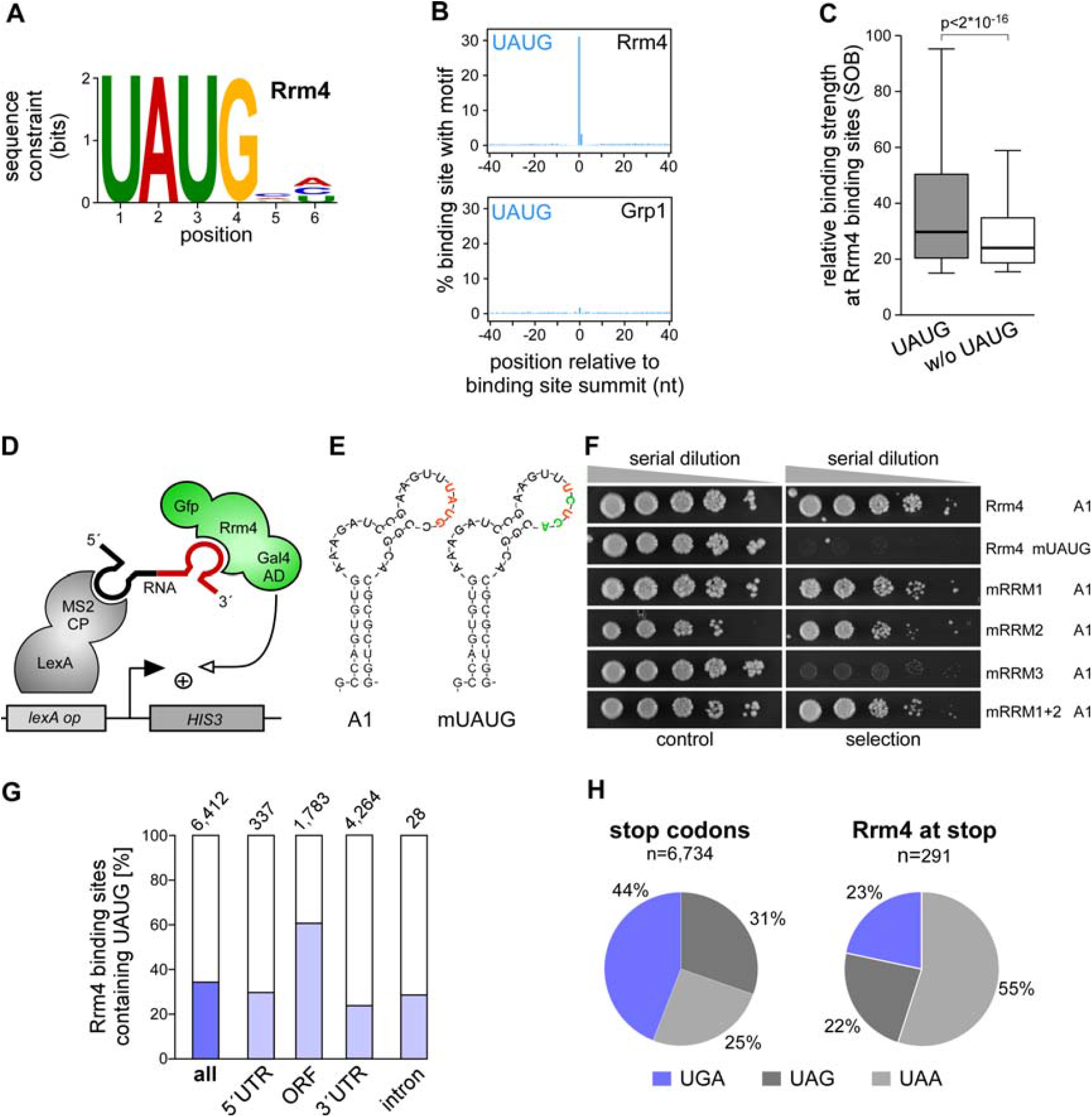
Rrm4 recognises UAUG *in vivo*. (**A**) Logo representation of the most enriched sequence motif at Rrm4 binding sites. At each position, the height of the stack is proportional to the information content, while the relative height of each nucleotide within the stack represents its relative frequency at this position. (**B**) Frequency of the Rrm4 motif UAUG around Rrm4 and Grp1 binding sites. Shown is the percentage of binding sites that harbour an UAUG starting at a given position in an 81-nt window around the binding site summit. (**C**) Box plot comparing relative binding strength at Rrm4 binding sites (signal-overbackground, SOB; see Materials and methods) with or without UAUG (n=2,201 and n=4,211, respectively; unpaired Student’s t-test). Box limits represent quartiles, centre lines denote 50th percentiles, and whiskers extend to most extreme values within 1.5x interquartile range. (**D**) Schematic representation of the yeast three-hybrid system: *lexA* operator (*lexA op*) sequences are bound by the LexA-MS2 coat protein (CP) hybrid (grey), recruiting the MS2-SELEX-RNA hybrid (black and red, respectively) to the promoter region of the *HIS3* reporter gene. Transcription is activated by binding of the third hybrid AD-Rrm4-Gfp (green) carrying a Gal4 activation domain (AD). (**E**) RNA structure prediction of aptamer SELEX-A1 with UAUG (red) or the mutated version mUAUG containing UCUC(A) (mutated bases in green). (**F**) Colony growth on control and selection plates of yeast cells expressing protein and RNA hybrids indicated on the right. RNA binding is scored by growth on selection plates (SC -his +3-AT, 3-amino-1, 2,4-triazole). mRRMx, Rrm4 variants harbouring mutations in RRM1, 2, 3 or 1 and 2. (**G**) Percentage of Rrm4 bs containing the motif UAUG, shown for all bs and separated into transcript regions. Total number of binding sites is indicated on top. (**H**) Relative contribution of the three stop codon variants to all (left) and Rrm4-bound stop codons (left). Although opal stop codons (UGA) fit best with a UAUG-containing binding site (UAUGA; present at 32 out of 63 bound UGA stop codons, 51%), they are depleted from Rrm4-bound stop codons.

A similar sequence analysis of the Grp1 binding sites initially suggested the sequence UGUA as a potential recognition motif (Fig EV5A). However, the same motif also frequently occurred at Rrm4 binding sites and showed no clear positioning relative to the Grp1 binding sites (Fig EV5B), making it questionable whether it was directly involved in the RNA recognition of Grp1. We therefore did not pursue this motif further.

In order to independently test whether Rrm4 specifically recognises the sequence motif UAUG, we applied the yeast three-hybrid assay (Fig 5D). We previously used this approach to successfully identify SELEX-derived RNA aptamers that were recognised by the third RRM domain of Rrm4 *in vivo* (RRM3) [36]. Intriguingly, two RNA aptamers, SELEX-A1 and SELEX-B1, contained the UAUG motif. We chose SELEX-A1 (Fig 5E) [36] to mutate the UAUG motif and tested RNA binding using the yeast three-hybrid assay. In contrast to the initial SELEX-A1 aptamer, the mutant version was no longer recognised by Rrm4 (Fig 5F; Fig EV5C). Consistent with earlier results, mutating the third RRM domain of Rrm4 gave similar results in the context of the initial SELEX-A1 aptamer [36]. Thus, our computational and experimental analyses indicate that Rrm4 specifically recognises the sequence motif UAUG via its third RRM domain.

Interestingly, Rrm4 binding sites in the whole ORF region showed a strong enrichment for the Rrm4 motif UAUG (Fig 5G), such that 61% of all ORF binding sites harboured UAUG. Since the start codon AUG is contained within this motif, the vast majority of start codon-associated binding sites exhibited the UAUG motif (88%, Fig EV5D). However, the frequency of Rrm4 binding was not increased compared to UAUG motifs in the surrounding sequence, indicating that Rrm4 does not show a particular preference for UAUG motifs in the context of translational start sites (Fig EV5E). Nevertheless, the Rrm4 recognition motif intrinsically overlaps with the start codon, suggesting a potential link to translation regulation. Binding of Rrm4 at the start codon might interfere, for example, with translational initiation of the bound target mRNAs.

In contrast to the UAUG prevalence in the ORF, stop codons seem to be recognised differently (UGA overlaps with UAUG i.e. UA<u>UGA</u>; but UAA was the most common stop codon, 55%, Fig 5H). Since UAUG-containing binding sites showed particularly strong Rrm4 binding (Fig 3C; as example, see *cdc12* in Fig 3E), Rrm4 appears to exhibit a tight association with the ORF via its third RRM domain. Uniting these observations, we hypothesised that Rrm4 simultaneously recognised multiple regions of the same cargo mRNAs. In line with this notion, we found that transcripts with a Rrm4 binding site in the 3’ UTR were significantly enriched for a second Rrm4 binding site in the ORF (663 out of 1,703 transcripts with at least two Rrm4 binding sites; p value < 2.22e-16, Fisher’s exact test). In 69% of these cases, the ORF binding site harboured UAUG, and in 56%, the 3’ UTR binding site of Rrm4 overlapped with a Grp1 binding site. Taken together, these observations would be consistent with a model that Rrm4 binds with its RRM domains RRM1 and/or RRM2 close to Grp1 in the 3’ UTR and via its third RRM domain to a UAUG-containing binding site in the ORF (Fig 6).

**Fig 6.**
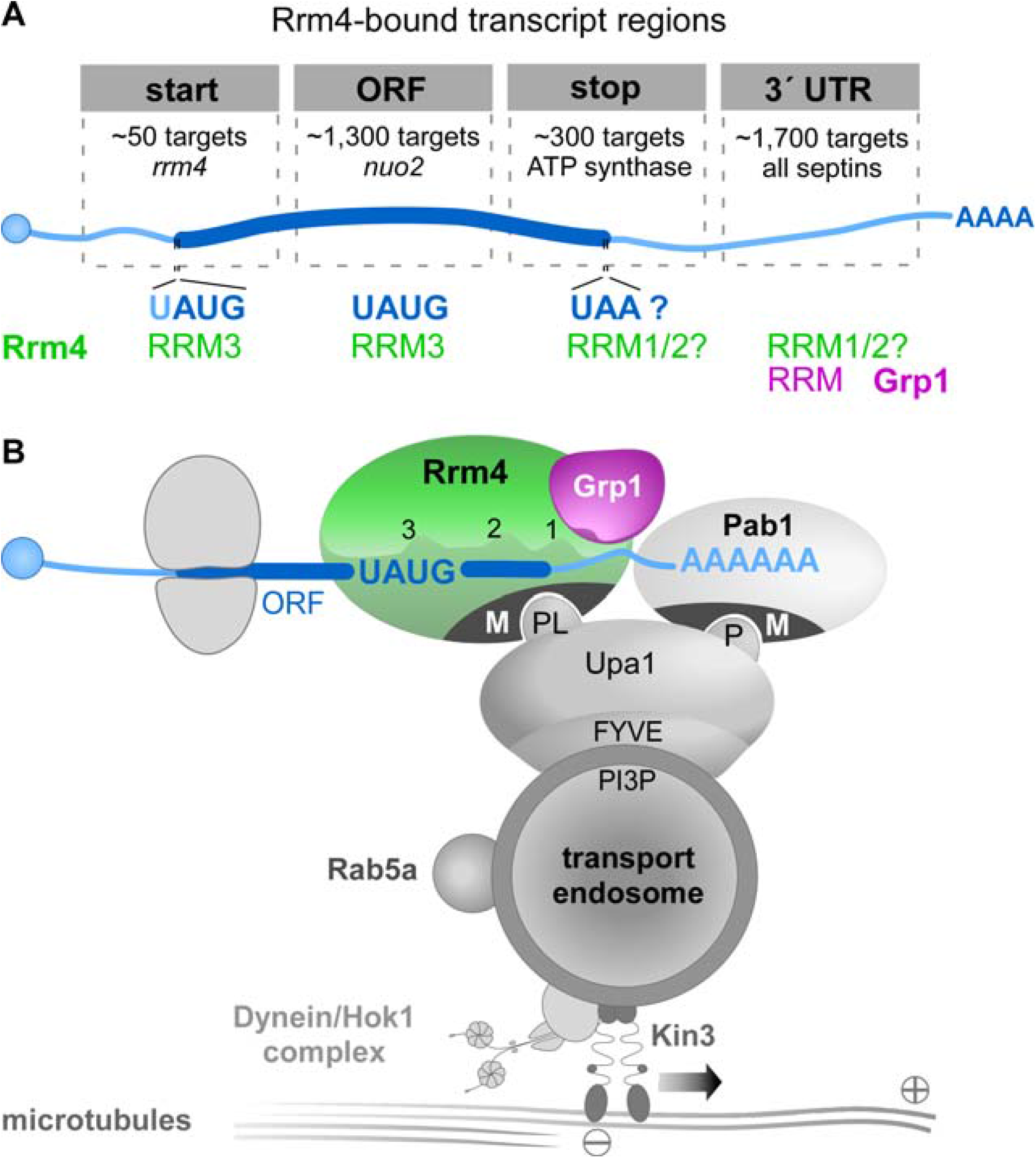
Model of Rrm4/Grp1-mediated endosomal mRNA transport. (**A**) Schematic drawing of target transcript with Rrm4-bound regions on top (5’ cap structure, blue circle; 5’ and 3’ UTR, light blue; ORF, dark blue; poly(A) tail, AAAA). Different categories of Rrm4 target transcripts were defined according to the presence of Rrm4 binding sites at the start codon, within the ORF, at the stop codon and in the 3’ UTR. Approximate number of target transcripts and selected examples are given for each category. About 900 of 1,300 target transcripts with an Rrm4 binding site in the ORF harbour an UAUG motif within the ORF binding site. Potential RRM domains of Rrm4 and Grp1 that may mediate RNA binding in the different transcript regions are given in green and magenta, respectively. (**B**) Simplified model proposing the spatial arrangement of endosomal RBPs with bound target transcripts. The three RRM domains of Rrm4 are schematically displayed and labelled by numbers (FYVE zinc finger domain; PI3P, phosphatidylinositol 3-phosphate; M, MademoiseLLE domain; P and PL, PAM2 and PAM2-like sequence, respectively; further details, see text). Note that complete endosomal mRNPs will consist of additional mRNA and protein components forming more complex, higher-order structures.

## Discussion

At present, a small number of high-throughput studies provide a global view on transported and localised mRNAs. In oocytes and embryos from fruit fly, transcriptome-wide RNA *in situ* hybridisation approaches revealed that the majority of transcripts exhibit a defined localisation pattern [37, 38]. In neuronal cells, the localised transcriptome and proteome have also been compiled [39, 40]. However, it is still unclear (i) how these cargo mRNAs reach their destination, (ii) which RBPs mediate their transport, and (iii) what are the precise interaction sites within the target mRNAs. Here, we applied iCLIP to study the newly identified endosomal RBP Grp1 and the key transport RBP Rrm4 during endosomal mRNP transport in *U. maydis*. To the best of our knowledge, this is the first detailed iCLIP analysis of mRNA transport.

### The GQ-rich RNA-binding protein Grp1 is a novel component of endosomal mRNPs

The small GQ-rich protein Grp1 shares similarity with other glycine-rich RBPs from humans and plants. A characteristic feature of this conserved class of RBPs is an N-terminal RRM domain followed by a glycine-rich low-complexity sequence that is typical for intrinsically disordered regions (IDRs). In RBPs, IDRs can mediate the assembly of membrane-less organelles through phase transitions [41], and this assembly is important, for example, during splicing [42]. Interestingly, IDRs have also been implicated in the formation of RNA granules for neuronal RNA transport [43, 44].

Small glycine-rich RBPs function in a wide variety of biological processes. The plant protein *At*GRP7, for example, is involved in cold stress adaption, osmotic stress response, circadian rhythm and plant immunity [21, 45, 46]. Human CIRBP regulates telomerase activity and the human cold shock protein RBM3 is involved in translational reprogramming during the cooling response of neuronal cells [20, 47, 48]. Globally, these proteins might function as RNA chaperones that prevent the formation of aberrant RNA secondary structures under stress conditions [45, 49].

In this study, we observe that loss of the fungal orthologue Grp1 causes aberrant alterations of the hyphal growth programme. Under optimal conditions *grp1Δ* hyphae grow significantly longer, suggesting an unusual acceleration of cell wall formation. Moreover, cell wall stress revealed clear abnormal morphologies in comparison to wild type. Consistent with a role in hyphal growth, Grp1 shuttles on Rrm4-positive endosomes that are the main transport units for long-distance transport of mRNAs in hyphae. Moreover, Grp1 and Rrm4 conjointly bind in the 3’ UTRs of thousands of target mRNAs (Fig 6A, see below). We therefore propose that the potential RNA chaperone Grp1 most likely constitutes an accessory component of endosomal mRNPs. Its function could be particularly important under suboptimal conditions. Alternatively, Grp1 might regulate stability and/or translation of mRNAs encoding proteins involved in hyphal growth independent of endosomal mRNA transport.

It is of note that the RRM protein Hrb27C (Hrp48) from fruit fly with a related domain architecture containing two N-terminal RRMs followed by a C-terminal GQ-rich region was found as an mRNP component during transport of *oskar* and *gurken* mRNAs [50-52]. Hence, the presence of small glycine-rich RBPs in transport mRNPs might be preserved across organisms.

### Endosomal RBPs recognise a broad spectrum of cargo mRNAs

In order to obtain a comprehensive view on the *in vivo* mRNA targets of an RBP, UV crosslinking techniques are currently the method of choice [53-55]. Here, we applied iCLIP to study fungal mRNA transport. A strength of our approach was the use of strains expressing functional Gfp-tagged versions of Grp1 and Rrm4 using homologous recombination to avoid overexpression artefacts. For application in fungi, we had to improve several steps [30], of which optimising the dose and duration of UV-C irradiation was most critical. Thereby, we were able to obtain a transcriptome-wide view of the cargo mRNAs, including their interaction sites with cognate RBPs at single nucleotide resolution.

Comparing two distinct RBPs present in endosomal mRNPs enabled us to disentangle the precise binding behaviour of the two co-localising RBPs. We identified more than 2,000 shared target transcripts, covering a substantial amount of the approximately 6,700 annotated protein-coding genes (http://pedant.helmholtz-muenchen.de/index.jsp; ORF update 2016.11.08) [31]. The broad target spectrum of endosomal mRNA transport fits with earlier observations that Rrm4 transported all mRNAs under investigation, albeit with different processivity of transport [17]. Moreover, loss of Rrm4 impairs the global mRNA distribution in hyphae, indicated by a disturbed subcellular distribution of the poly(A)-binding protein Pab1 [17]. Thus, one function of the endosomal mRNA transport machinery might be the equal distribution of mRNPs to supply all parts of the hyphae with mRNAs. This might be particularly important for those parts that are distant from the mRNA-synthesising nucleus. Such a universal mRNA transport mode resembles the “sushi belt model” in neuronal cells, in which the shuttling of mRNPs by active transport along microtubules is thought to distribute mRNAs throughout the dendrite to serve synapses that are in demand of mRNAs [56]. Furthermore, since many mRNAs in *U. maydis* appear to be transported in a translationally active state (see below), this mRNA distributer function would also disclose the mechanism of how ribosomes are transported by endosomes, as observed previously [57].

Notably, we found that most nuclear-encoded subunits and accessory components of mitochondrial F_O_F_1_ ATPase are targets of endosomal mRNP transport in *U. maydis*. Consistent with the idea that a precise spatiotemporal regulation of translation might be important for efficient mitochondrial protein import, we observed that the abundance of Atp4 is reduced in *rrm4Δ* hyphae [18]. These results agree with previous findings that the 3’ UTR of *ATP2* mRNA from *S. cerevisiae* is important for efficient mitochondrial uptake of mature Atp2p [58]. A close link between RNA biology and mitochondrial protein import is thus an emerging theme in mitochondrial biology [59-63].

### Distinct binding patterns of Rrm4 may allow orchestration of mRNA transport and translation

The positional information obtained by single nucleotide resolution was essential to uncover the precise binding behaviour of the involved RBPs. In the majority of cases, Rrm4 binds together with Grp1 in the 3’ UTR (1,700 transcripts; Fig 6A). The vicinity of these binding sites to the poly(A) tail fits the previous observation that the endosomal adaptor protein Upa1 interacts with both Rrm4 and Pab1 [64]. While translating ribosomes would potentially remove RBPs from the ORF [65], such binding in the 3’ UTR, as seen e.g. on all four *septin* mRNAs, would allow simultaneous translation and transport of mRNAs. Consistently, we have recently provided evidence that endosome-coupled translation of *septin* mRNAs mediates endosomal assembly and transport of heteromeric septin complexes [6, 19]. Since transport of translationally active mRNAs has recently been observed in neurons [66], this mode of transport might be more widespread than currently anticipated.

Our transcriptome-wide RNA binding maps illustrate an intriguing binding pattern of Rrm4 at translational landmark sites, indicating an intimate connection of endosomal mRNP transport and translation in *U. maydis* (Fig 6A): First, in over 1,000 target transcripts Rrm4 binds within the ORF, indicating that it may be involved in stalling translational elongation. Notably, a similar mechanism was suggested in neurons, in which a subset of mRNAs are translationally stalled during transport [67]. Similar to Rrm4, the neuronal RBP FMR1 binds its translationally stalled target mRNAs preferably in the coding sequence [68]. Second, in about 50 and 300 target transcripts, respectively, Rrm4 precisely binds at the start and stop codons, suggesting modulation of translation initiation and termination. At start codons and within the ORF, the Rrm4 binding sites frequently harboured UAUG. This motif is recognised by the third RRM domain of Rrm4. In accordance with its ELAV-type domain organisation, we therefore propose that Rrm4 binds UAUG-containing binding sites via its third RRM to influence translation (Fig 6B), while the two tandem RRMs (RRM1/2) bind the target mRNAs in the 3’ UTR, possibly together with Grp1 (Fig 6B). In a previous study, we observed that mutations in RRM1 and RRM3 led to strongly reduced overall RNA binding of Rrm4, although mutations in RRM3 did not show a mutant phenotype with respect to hyphal growth [16]. In contrast, mutations in RRM1 affected hyphal growth strongly [16], accompanied by reduced endosomal shuttling of Pab1 as well as *cdc3* mRNA [6]. Therefore, the potential translational regulation during endosomal mRNP transport mediated by RRM3 may be an additive and used for fine-tuning. The tandem RRM domains RRM1 and RRM2 of Rrm4 might mediate the recognition of target mRNAs for transport, possibly via binding in the 3’ UTR.

In essence, our comprehensive transcriptome-wide view on endosomal mRNA transport revealed the precise positional deposition of co-localising RBPs during transport. The key RBP Rrm4 exhibits a particular binding behaviour by recognising distinct landmarks of translation in target mRNAs. Thereby, translation and transport might be intimately coupled and precisely synchronised for the specific expression needs of each target transcript.

## Materials and methods

### Plasmids, strains and growth conditions

Cloning was done using *E. coli* K-12 derivate Top10 (Life Technologies, Carlsbad, CA, USA). All strains were constructed by the transformation of cells with linearised plasmids, and homologous integration events were verified by Southern blot analysis [69]. Genomic DNA of wild type strain UM521 (*a1b1*) served as template for PCR amplifications unless noted otherwise. Proteins were tagged with eGfp (enhanced green fluorescent protein; Clontech, Mountain View, CA, USA), Tag-Rfp or mCherry [26, 70]. Plasmid sequences are available upon request. The accession numbers of *U. maydis* genes used in this study are: *rrm4* (UMAG_10836), *grp1* (UMAG_02412), *pab1* (UMAG_03494). Detailed information is supplied in Appendix Tables S1-S3. The conditions used for cultivation of *U. maydis* are described elsewhere [69]. For the induction of filamentous growth of lab strain AB33 and derivates, cultures were grown to an OD600 of 0.5 in complete medium supplemented with 1% glucose (CM-glc) and shifted to nitrate minimal medium supplemented with 1% glc (NM-glc). The growth curves of *U. maydis* strains were recorded by cultivating strains in CM-glc at 28°C and measuring the OD_600_ every 2 h.

### Multiple sequence alignments

Orthologous proteins were identified using BLAST P (https://blast.ncbi.nlm.nih.gov/blast/). Clustal W and GeneDoc 2.6 were used for multiple amino acid sequence alignment and graphical representation, respectively [71].

### Preliminary affinity purification experiments using Rrm4-GfpTT as bait

Preliminary purification experiments were performed using a strain expressing Rrm4 fused C-terminally to Gfp and tandem affinity purification (tap) tag [16]. We analysed the eluted fractions by mass spectrometry either as complex mixture in solution (Fig EV1C) or by cutting individual bands from an SDS-PAGE gel (Fig EV1D). In both cases, the protein purification followed a published protocol (CLIP) [16]. Both experiments were considered to be pilot studies and were only performed once. In case of the in-solution digest, the protein samples were precipitated with 20% trichloroacetic acid (1:2, v/v) and washed five times with cold acetone. Samples were digested with sequencing-grade modified trypsin (Promega), and the resulting peptides were separated into fractions by nanoLC (PepMap100 C-18 RP nanocolumn and UltiMate 3000 liquid chromatography system, Dionex). Fractions were spotted with matrix solution on a MALDI plate and subsequently analysed by MALDI-TOF MS (4800 Proteomics Analyser, AB Sciex). Mass spectrometry data were searched against the MUMDB (Munich *Ustilago Maydis* Data Base; 6785 entries; IBIS Institute of Bioinformatics and Systems, German Research Center for Environmental Health) [31] using Mascot embedded into GPS Explorer software (AB Sciex). The total peptide score is the sum of all peptide scores corresponding to the predicted proteins, excluding the scores of duplicate matches. The best peptide score is the best score from all identified peptides corresponding to the predicted proteins.

Applying as filter a total peptide score of larger than 46, we identified 23 proteins as potential interaction partners of Rrm4 (Fig EV1C). Among these proteins were interesting candidates that were identified before in independent approaches: Chitinase Cts1 and ribosomal protein Rps19 were found in a differential proteomics study, comparing wild type and *rrm4Δ* hyphae [18]. Upa2 is a PAM2 motif-containing protein that was identified as potential interaction partner of Rrm4 in a bioinformatics approach [64]. Grp1 (UMAG_02412) was chosen for further analysis, since it contained an RNA recognition motif (RRM) for RNA binding. In the gel-based approach, the calmodulin beads were washed three times and incubated in SDS-loading buffer for 8 min at 90°C. The supernatant was subjected to SDS-PAGE (10% polyacryl amide). Proteins were staining with Coomassie Blue. After excision of selected bands samples were destained and dried under vacuum. The dried material was suspended in 30 μl 10 mM ammonium bicarbonate (pH 8.3) containing 0.6 μg of trypsin (Promega) and 10% acetonitrile. After incubation at room temperature for 12 h, the trypsin digest was extracted and analysed as described above. We were able to verify Rrm4-GfpTT (TEV cleavage product 117 kDa) and Grp1 (18 kDa; Fig EV1D) in selected bands matching their expected gel position after electrophoresis.

### Colony morphology, temperature stress and cell wall stress

For growth on solid media, cell suspensions (OD600 of 0.5) were inoculated onto respective plates at 28°C. Colony morphology was tested on CM (1% glc) medium. Temperature stress was analysed on CM (1% glc) medium, incubated at 28°C, 20°C or 16°C. Cell-wall stress was analysed by supplementing CM (1% glc) medium with 50 μM Calcofluor White (Sigma-Aldrich, Taufkirchen, Germany) or 40 μg/ml Congo Red (Sigma-Aldrich). All plates were kept for at least 48 h at the required temperature. The set-up used to acquire images was described before [64]. To analyse cell wall stress in hyphae, 2.5 μM Calcofluor White was added to cultures 3 hours post induction (h.p.i) and incubated for an additional 2 h.

### Microscopy and image processing

The microscope set-ups and dual-colour imaging were used as described before [5, 25, 72]. Gfp and tagRfp or mCherry fluorescence was simultaneously detected using a two-channel imager (DV2, Photometrics, Tucson, AZ, USA). All images and videos were acquired and analysed using Metamorph (Versions 7.7.0.0 and 7.7.4.0; Molecular Devices, Sunnyvale, CA, USA).

Cell length was assessed by measuring the length of single cells from pole-to-pole. For hyphae, empty sections were not included in the measurements. The length of empty sections was assessed by measuring the distance from septum to septum of the first empty section at the distal pole of hyphae. Cell wall defects induced by Calcoflour White were quantified by manually scoring for the presence of abnormal cell wall shapes.

For the analysis of co-localisation and velocity of moving signals the acquired videos were converted to kymographs using Metamorph. Co-localisation was assessed by quantifying kymographs acquired by dual-colour imaging. Changes in direction were counted as individual signals. Processive signals (distance travelled > 5 μm) were counted manually. Velocity was only measured for processive signals (movement > 5 μm). For all quantifications, at least three independent experiments were analysed. Statistical analysis was performed using Prism5 (Graphpad, La Jolla, CA, USA).

The gradient of Grp1-Gfp, Pab1-Gfp and Gfp in the presence and absence of Rrm4 was quantified by measuring the fluorescence intensity in a previously specified region of interest (ROI; ROI1 in vicinity to the nucleus and ROI2 at the hyphal tip). The fluorescence close to the nucleus was then set in relation to the signal intensity at the tip. Statistical analysis was performed using Prism5 (Graphpad, La Jolla, CA, USA).

### iCLIP experiments

The iCLIP protocol was modified from [30] and the original application in *U. maydis* [17]. (i) The initial two-step tap tag purification [17] was switched to single-step purification using strains expressing C-terminal Gfp fusions and Gfp-trap immunoprecipitation (GFP-Trap®_MA, ChromoTek, Martinsried, Germany) [73]. (ii) For UV-C crosslinking, cells were irradiated continuously in a single session. (iii) The addition of DNase and RNase I was omitted from the samples used for iCLIP library preparation. Due to the high RNase activity in fungal cells, the addition of external RNases was unnecessary, therefore RNase I (10 Units/sample; 16 minutes, 37°C; Thermofisher Scientific, Darmstadt, Germany) was only added in the control experiments (Fig 3A).

All iCLIP experiments were performed with 150 ml cultures grown to an OD600 = 0.5 in CM-medium (supplemented with 1% glc) and then shifted to NM-medium (supplemented with 1% glc) to induce hyphal growth. After 6 h, each culture was split into 3 x 50 ml, harvested by centrifugation at 6,280 g for 10 min at 4°C, and the cell pellets were resuspended in 5 ml ice-cold PBS. For UV-C crosslinking, cells were irradiated once with 200 mJ/cm^2^ at 254 nm (Biolink UV-Crosslinker, Vilber-Lourmat, Eberhardzell, Germany) as a thin layer in a square petri dish (10 cm^2^), pooled in a 50 ml tube, and harvested by centrifugation at 6,280 g for 10 min at 4°C. The cell pellet was resuspended in 6 ml lysis buffer (50 mM Tris-HCl, pH 7.4, 100 mM NaCl, 1% Nonidet P-40, 0.1% SDS, 0.5% sodium deoxycholate) supplemented with inhibitors (per 10 ml lysis buffer): 2× Complete protease inhibitor EDTA-free (Roche Diagnostics, Mannheim, Germany), 1 mM reduced DTT (GERBU, Heidelberg, Germany), 5 mM Benzamidine (Sigma), 1 mM PMSF (Sigma), 0.75 μg/μl heparin (Sigma), 5.25 ng/μl pepstatin A (Sigma) and 15 μl SUPERase-In (20 U/μl; Thermo Fisher Scientific). The solution was split into three times 2 ml for cell lysis and immunoprecipitation. Cell lysis was performed in a Retsch ball mill (2 ml cell suspension/grinding jar, 2 grinding balls, d=12 mm; MM400; Retsch, Haan, Germany) 3 x 10 min at 30 Hz while keeping samples frozen using liquid nitrogen. All of the following steps up to the 5’ labelling of RNA were performed at 4°C. The cell lysates were pooled in a 15 ml tube, split in precooled 1.5 ml tubes and cleared by centrifugation at 16,200 g for 15 min at 4°C. The protein-RNA complexes were immunoprecipitated from each strain by using a total of 60 μl GFP-Trap magnetic agarose beads (Chromotek; GFP-Trap^®^_MA) [73]. Before immunoprecipitation, the 60 μl beads were added to a 2 ml tube and pre-washed three times with 500 μl ice-cold lysis buffer (without inhibitors) and afterwards split into three times 20 μl beads in 2 ml tubes. For immunoprecipitation, 2 ml cleared cell lysate were added to 20 μl beads and the tubes were incubated rotating for 1 h at 4°C. Next, the beads were combined in a 2 ml tube and washed three times with 900 μl high-salt wash buffer (50 mM Tris-HCl, pH 7.4, 1 M NaCl, 1 mM EDTA, 1% Igepal CA-630, 0.1% SDS, 0.5% sodium deoxycholate) and four times with 900 μl PNK wash buffer (20 mM Tris-HCl, pH 7.4, 10 mM MgCl_2_, 0.2% Tween-20). For each subsequent enzymatic reaction, the 60 μl beads were split into three times 20 μl beads in 2 ml tubes and re-pooled for the washing steps. 3’ end RNA dephosphorylation, L3 adapter ligation, 5’ end phosphorylation, SDS-PAGE and nitrocellulose transfer were performed as described [30] with minor changes implemented. For 5’ end phosphorylation, 50% of each sample was radioactively labelled with ^32^P-γ-ATP for 10 min at 37°C and the labelled beads were washed once with PNK wash buffer before they were combined with the unlabelled beads. The beads were diluted in 80 μl NuPAGE LDS loading buffer (Thermo Fisher Scientific) with 0.1 M DTT added. Samples were heated at 70°C for 5 min and split among two adjacent wells on a 4-12% NuPAGE Bis-Tris gel (Thermo Fisher Scientific). The protein-RNA complexes were separated at 180 V for 70 min in MOPS running buffer with NuPAGE reducing agent added (Thermo Fisher Scientific) and transferred onto a nitrocellulose membrane. For Western blotting, Gfp was detected using monoclonal α-GFP primary antibodies (clones 7.1 and 13.1; Sigma) and a mouse IgG HRP conjugate (H+L; Promega, Madison, WI, USA) as secondary antibody. Peroxidase activity was determined using the AceGlow blotting detection system (Peqlab, Erlangen, Germany).

Labelled RNA was detected by exposure of X-ray films ranging from 2 h - overnight at −80°C. cDNA library preparation was performed as described before [30). To avoid over-amplification of the cDNA library, the optimal number of PCR cycles producing PCR products of around 150 nt (cDNA insert = 20-30 nt; L3 adapter, RT-primer and P3/P5 Solexa primers = 128 nt) was tested for each protein in every experiment (PCR cycler PTC-200, MJ Research, St. Bruno, Quebec, Canada; Fig EV2D). For all Grp1 and Gfp replicates, 18 PCR cycles were determined to be optimal, while the optimal PCR cycle numbers for the Rrm4 replicates were determined to be 18 and 22 PCR cycles. The iCLIP libraries were multiplexed and sequenced on an Illumina HiSeq 2500 (San Diego, CA, USA; 51-nt reads, single-end), yielding a total of 118 million reads.

### iCLIP data processing

All bioinformatics analyses are based on the *U. maydis* 521 genome sequence (original PEDANT database name p3_t237631_Ust_maydi_v2GB) and the associated gene annotation (version p3_t237631_Ust_maydi_v2GB.gff3; both downloaded from ftp://ftpmips.gsf.de/fungi/Ustilaginaceae/Ustilago_maydis_521/). We extended all genes by 300 nt on either side to include potential 5’ and 3’ UTR regions which are currently not annotated in the *U. maydis* genome. For manual annotation of transcript ends, RNASeq data (AB33 hyphae, 6 h.p.i.) were used, and transcript ends were defined at the position where read coverage dropped below 10.

Basic quality checks were applied to all sequenced reads using FastQC (https://www.bioinformatics.babraham.ac.uk/projects/fastqc/). Afterwards, iCLIP reads were filtered based on sequencing quality (Phred score) in the barcode region, keeping only reads with at most one position with a Phred score < 20 in the experimental barcode (positions 4 to 7) and without any position with a Phred score < 17 in the random barcode (positions 1 to 3 and 8 to 9). The reads were then de-multiplexed based on the experimental barcode at positions 4 to 7 using Flexbar (version 2.4, GitHub, San Francisco, CA, USA) without allowing mismatches [74). The following analysis steps were applied to all individual samples: remaining adapter sequences were trimmed from the 3’ end of the reads using Flexbar (version 2.4) allowing one mismatch in 10 nt, requiring a minimal overlap of 1 nt between read and adapter as well as removing all reads with a remaining length of less than 24 nt (including the 9-nt barcode). The first 9-nt of each read containing the barcode were trimmed off and added to the read name in the fastq file.

Filtered and trimmed reads were mapped to the *U. maydis* genome and its gene annotation using STAR (version 2.5.1b, GitHub) [75], allowing up to two mismatches and without soft-clipping on the 5’ end of the reads. Only uniquely mapped reads were kept for further analysis.

After mapping and filtering, duplicate reads were marked using the *dedup* function from bamUtil (version 1.0.7; https://github.com/statgen/bamUtil) and removed if carrying an identical random barcode, and hence representing technical duplicates. The nucleotide position upstream of each aligned read was considered as the ‘crosslink nucleotide’, with each read counted as individual ‘crosslink event’. The total number of crosslink events for the different iCLIP libraries can be found in Fig EV3A. To assess the reproducibility between biological replicates (Fig EV3C), we counted the number of crosslink events within each gene.

We then proceeded to define putative RBP ‘binding sites’, i.e. sites that show a significant enrichment of crosslink events compared to the surrounding sequence. Since most RBPs contact multiple positions in the RNA, the crosslink events in a binding site commonly spread across more than one nucleotide. For the identification of such windows with significantly increased crosslink event frequency, peak calling was performed using ASPeak that accounts for the different expression levels of the underlying transcripts [76]. Peak calling was performed on merged replicates for each RBP to maximise the signal. In order to obtain binding sites of a uniform width, we further processed the predicted peaks [77]. We first determined the summit of each peak (i.e. the position with highest number of crosslink events within the peak; if two or more positions showed the same count, the most upstream position was taken) and then defined a binding site window of 9-nt that was centred around the summit position. Overlapping peaks were merged and newly centred on the summit of the combined window as described above. Finally, we filtered the binding sites as follows: (i) To account for reproducibility, we required each binding site to be detected by at least 5 crosslink events from each biological replicate. (ii) We further removed all Rrm4 and Grp1 binding sites that overlapped by at least 1 nt with any of 88 reproducible Gfp binding sites. To define the set of target genes that were exclusively bound by Rrm4 and showed no evidence of Grp1 binding (‘Rrm4-unique targets’), we subtracted the intermediate list of binding sites at this point for Grp1 from the final Rrm4 binding sites (after SOB filtering, see below), and vice versa.

For the remaining analysis, we included an additional processing step to focus on the top 25% of binding sites. To determine this set, we used the ratio of crosslink events within the binding site over the background crosslink events in the surrounding sequence (termed ‘signal-over-background’, SOB) as a proxy of binding site strength: Crosslink events from both replicates were summed up within each binding site and divided by the gene-specific background to obtain an SOB value for each binding site. The background for each gene was defined as the number of crosslink events outside the binding sites divided by the number of nucleotides in the gene that harbour such background crosslink events. We note that while this procedure alleviates the impact of transcript-level differences, it does not correct for UV crosslinking bias and similar confounding factors, and should therefore only be used as a proxy for bulk comparisons of binding sites. Only binding sites within the top 25% of the SOB distribution for each RBP were taken into consideration for further analyses.

This SOB filtering procedure yielded a total of 6,412 and 6,478 binding sites for Rrm4 and Grp1, respectively (Fig 3B, Fig EV3). The binding sites corresponded to 3,262 and 3,165 target transcripts for Rrm4 and Grp1, respectively. When assigning genomic nucleotides and RBP binding sites to distinct transcript regions (Fig 3D), we applied the following hierarchy to resolve overlapping annotation: 3’ UTR > 5’ UTR > exon > intron. Binding sites of Rrm4 and Grp1 were considered as overlapping if at least 1 nt was shared between the 9-nt binding sites.

To study the distribution of binding sites of Rrm4 and Grp1 in 3’ UTRs (Fig 4C), we manually annotated transcript ends. This was necessary, because the *U. maydis* gene annotation only includes ORF regions of genes. To this end, RNASeq data from AB33 hyphae (6 h.p.i.) were visually inspected and 3’ ends were defined due to strong reduction in RNASeq coverage.

Enriched sequence motifs around RBP binding sites were identified using DREME [35; parameter –norc to search the coding strand only) to analyse a 30-nt window around all binding site summits compared to shuffled nucleotides. Based on the sequence profile and the UAUG content around binding sites (Fig 5A-B), we counted a binding site as UAUG-containing if it harboured the motif with the last 5 nt of the 9-nt binding sites. In order to assess the frequency of Rrm4 binding sites at UAUG motifs at start codons and in other regions (Fig EV5E), we used all transcripts that carry Rrm4 binding sites anywhere within the transcript and are hence sufficiently expressed to be detected in our iCLIP analysis.

Analysing functional categories of cellular components on identified targets was performed using the FunCat annotation scheme (http://mips.gsf.de/funcatDB/; version 2.1, reference genome: p3_t237631_Ust_maydi_v2GB) [78]. Categories were filtered during analysis for enrichment by setting the p-value threshold to <0.05.

### RNASeq library preparation and data processing

RNA was extracted from AB33 hyphae 6 h.p.i. using the RNeasy Mini kit following the manufacturer’s instructions for preparation of total RNA from yeast (Qiagen, Hilden, Germany). To this end, AB33 hyphae were opened in a Retsch ball mill (3 balls, d=4 mm; MM400; Retsch, Haan, Germany) 4 times for 5 min at 30 Hz while keeping samples frozen using liquid nitrogen. The resulting cell powder was resolved in 450 μl RLT buffer (+β-mercaptoethanol) and centrifuged at 13,000 rpm for 2 min at 4°C. The supernatant was transferred to a new reaction tube, mixed with 1 volume 70% EtOH and then added to the RNeasy spin column. All following processing steps were performed according to manufacturer’s instructions. TruSeq RNA Library Prep kit v2 (Illumina, San Diego, CA, USA) was used for cDNA library generation. The cDNA libraries were sequenced using the HiSeq 2000 platform (Illumina) with 151-nt single-end reads.

Basic quality checks were applied to all sequenced reads using FastQC (https://www.bioinformatics.babraham.ac.uk/projects/fastqc/). Afterwards, RNASeq reads were trimmed based on sequencing quality (Phred score) using Flexbar (version 3.0.3) [74]. Specifically, adapter sequences were removed (TruSeq Universal Adapter), and reads were trimmed at the first position with a Phred score < 20 and removed if the remaining read length was less than 20 nt. The trimmed reads were mapped to the *U. maydis* genome and its gene annotation using STAR (version 2.5.3a) [75], allowing up to five mismatches with soft-clipping. Uniquely mapped reads were kept for further analysis.

### Yeast three-hybrid analysis

Yeast three-hybrid experiments were performed as described previously [36, 79]. To test the interaction with Rrm4, the plasmids (Appendix Table S4) encoding the RNA aptamers SELEX-A1[36] or mutated SELEX-A1 (mUAUG; this work) were cotransformed in strain L40 coat with the corresponding plasmids encoding for Rrm4 or mutated variants [36, 80]. Transformed cells were incubated on SC -ura -leu plates (2-3 d at 28°C) before single clones were selected. Interaction was assayed as growth on selection medium SC -his +1 mM 3-AT (3-amino-1,2,4-triazole; Sigma-Aldrich; 3 d at 28°C). For the serial dilution assays, single clones were grown in SC -ura -leu medium to a starting OD600 = 0,5 and sequentially diluted 1:5 in water. The dilutions were then spotted on control (SC -ura -leu) and selection (SC -his +1 mM 3-AT) plates and incubated at 28°C.

### Data Availability

The iCLIP and RNASeq dataset are available from GEO under the accession numbers GSE109557 and GSE109560, respectively. The associated SuperSeries is GSE109561.

## Supporting information

## Acknowledgements

We thank Dr. J. Kahnt for pilot Tap tag experiments and Dr T. Pohlmann for initial work on Grp1 fusion proteins. We acknowledge R.F.X. Sutandy for iCLIP support, M. Brüggemann for RNASeq data analysis, F. Finkernagel for bioinformatics and Dr. M. Seiler for critical reading of the manuscript. We are grateful to U. Gengenbacher and S. Esch for excellent technical assistance and members of the IMB Genomics core facility for technical assistance. The work was funded by grants from the Deutsche Forschungsgemeinschaft to MF (FE 448/8-1; FOR2333-TP03 FE448/10-1; CEPLAS EXC1028) and KZ (FOR2333-TP10 ZA881/1-1) and JK (SPP1935 KO4566/2-1) and JU (European Research Council; 617837-Translate).

## Author contributions

LO, CH, JU, JKö, MF and ZK designed this study and analysed the data. LO and CH performed experiments to characterise Grp1 and comparative iCLIP analysis. JKoe performed preliminary affinity purification experiments. SB, AB, and ZK performed all computational iCLIP data analyses. MF and ZK drafted and revised the manuscript with input from all co-authors. MF and ZK directed the project.

---

**Figure EV1.**
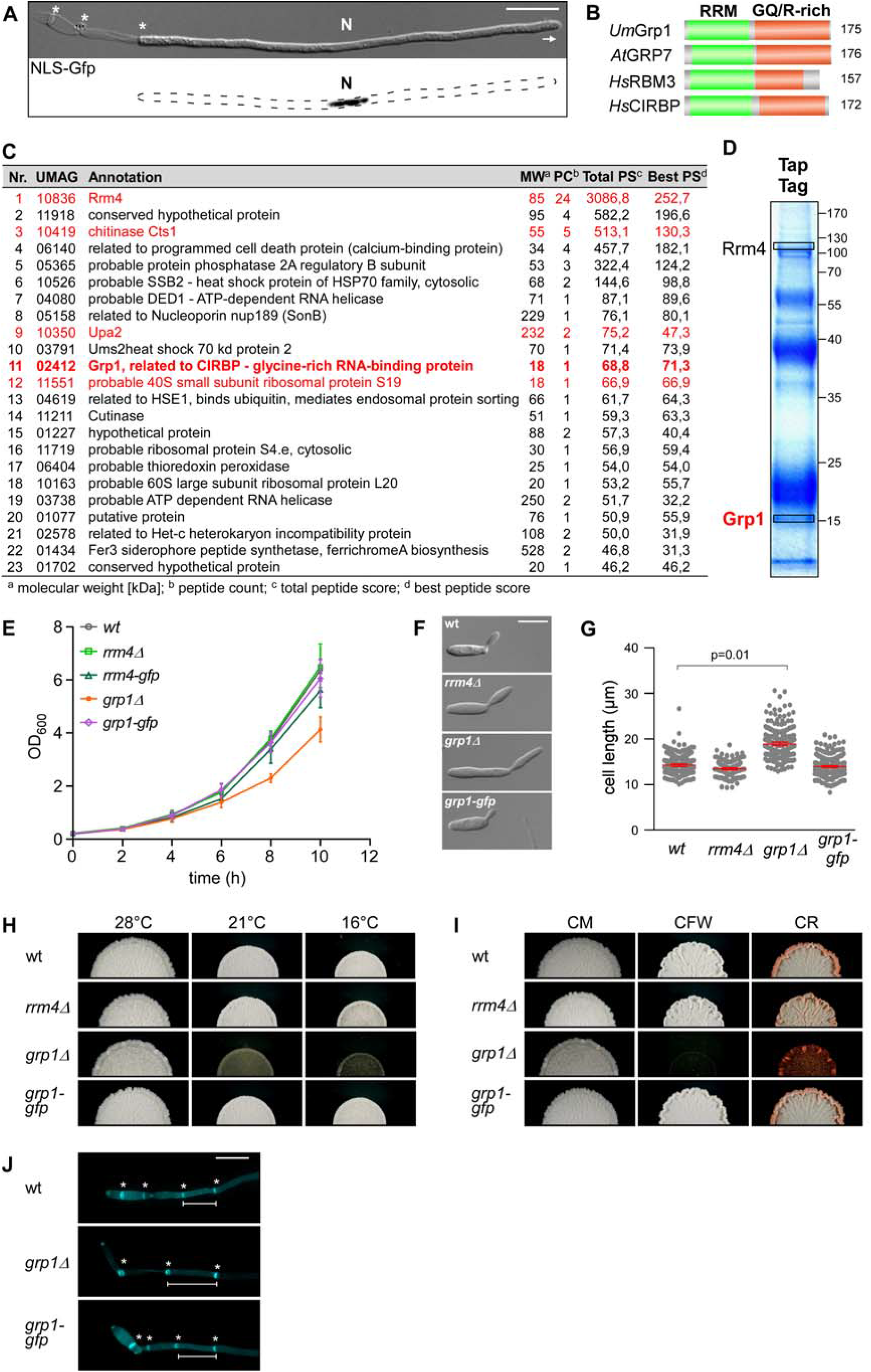
Loss of Grp1 causes defects in cell growth. (**A**) Hyphal form (6 hours post induction, h.p.i.) of laboratory strain AB33 expressing a Gfp-tagged protein with nuclear localisation signal to stain the nucleus (N; λN-NLS-Gfp, phage protein λN fused to triple Gfp, containing a nuclear localisation signal; inverted fluorescence image shown; scale bar, 10 μm). Hyphae expand at the apical pole (arrow) and insert septa (asterisks) at the basal pole in regular time intervals resulting in the formation of empty sections. (**B**) Schematic representation of the domain architecture of four small glycine-rich proteins (RRM, RNA recognition motif, green; GQ/R, glycine-rich region with arginine or glutamine, red). *Um*Grp1 from *U. maydis* (UMAG_02412), *At*GRP7 from *Arabidopsis thaliana* (RBG7; NC_003071.7), *Hs*RBM3 and *Hs*CIRBP from *Homo sapiens* (NC_000023.11 and NC_000019.10, respectively). Number of amino acids indicated on the right. (**C**) Results of preliminary affinity purification experiments using Rrm4-GfpTT as bait (see Materials and methods). Proteins with a functional link to Rrm4 are marked in red (this study) [18, 64]. Peptide count: number of identified peptides corresponding to predicted protein; total peptide score: sum of all peptide scores corresponding to predicted protein, excluding the scores of duplicate matches; best peptide score: best score from all identified peptides corresponding to predicted protein. Note that the difference between total peptide score and best peptide score is a correction of the software depending on how many possible predicted candidates match to the identified peptide mass. (**D**) Tandem affinity purification using Rrm4-GfpTT as bait. Protein bands were stained with Coomassie Blue after SDS-PAGE. Proteins in boxed areas were identified as Rrm4 and Grp1 (size of marker proteins in kDa on the right). (**E**) Growth curve of indicated AB33 derivatives growing in liquid culture. Data points represent averages from three independent experiments (n=3). Error bars show s.d. (**F**) Differential interference contrast (DIC) images of AB33 derivatives as yeast-like budding cells (scale bar, 10 μm). (**G**) Length of budding cells (shown are merged data from three independent experiments, n=3; > 100 cells per strain were analysed, *wt*, 269; *rrm4Δ*, 122 (only two independent experiments); *grp1Δ*, 263; *grp1G*, 318), overlaid with the mean of means, red line, and s.e.m.; paired two-tailed Student’s t-test on the mean cell lengths from the replicate experiments. (**H**) Colonies of indicated AB33 strains grown in the yeast form incubated at different temperatures (28°C for 1 d or 16°C and 21°C for 5 d). (**I**) Colonies of indicated AB33 strains grown in the yeast form. Incubated plates contained cell wall inhibitors (CM, complete medium for 1 d; CFW, 50 μM Calcofluor White for 4 d; CR, 57.4 μM Congo Red for 4 d). (**J**) Fluorescence images of the basal pole of hyphae of AB33 derivatives (6 h.p.i.). Septa (asterisks) were stained with CFW. White bars indicate exemplary length measurements of empty sections shown in Fig 1D (scale bar, 10 μm).

**Figure EV2.**
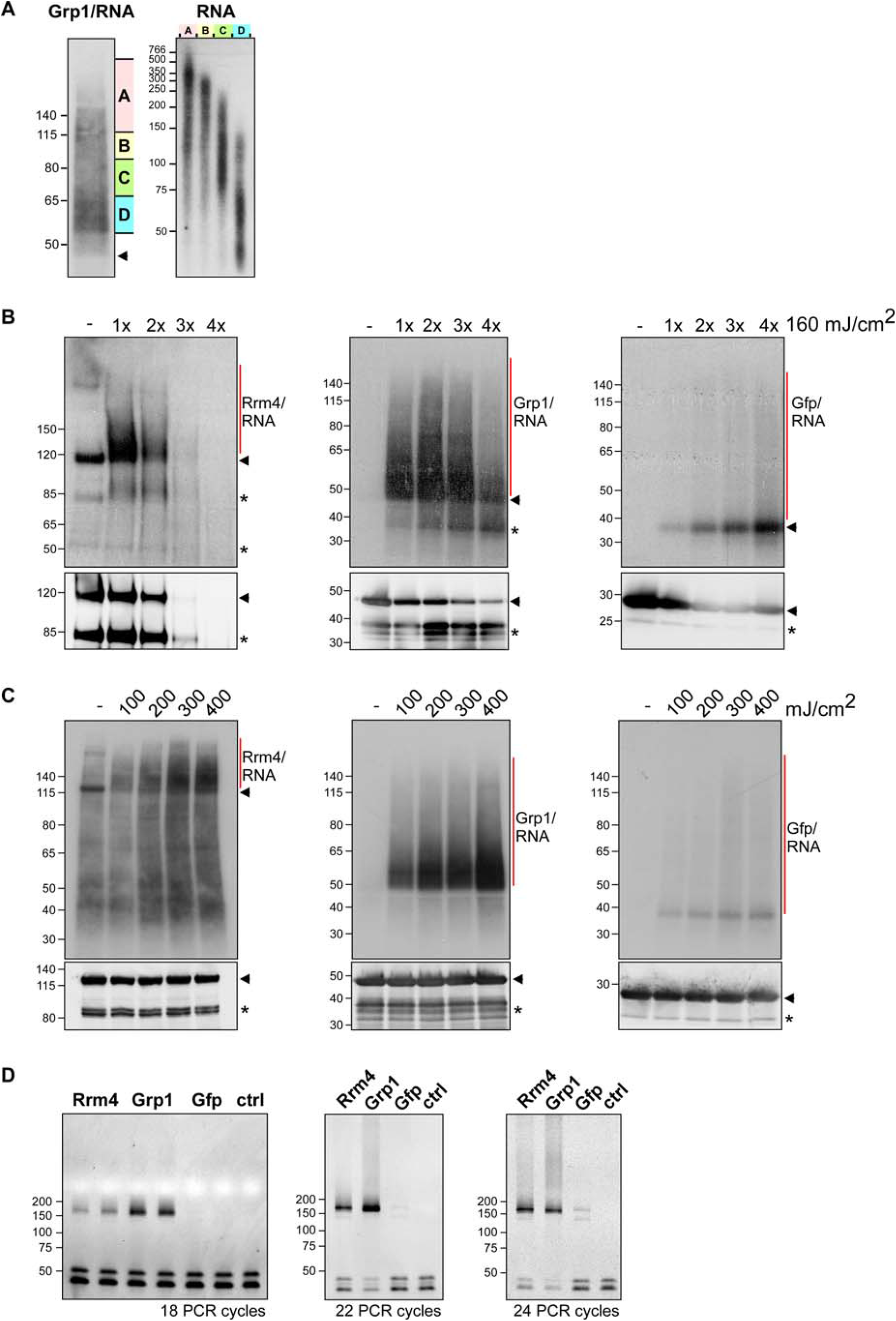
Improving the iCLIP protocol for fungal RBPs. (**A**) Grp1-Gfp/RNA complexes were size-separated on denaturing PAGE after UV-C irradiation and transferred to a nitrocellulose membrane (left). RNA was radioactively labelled, and protein-RNA complexes with covalently linked RNAs of different sizes were visible as smear above the expected molecular weight of the Grp1-Gfp protein (45 kDa; marked by arrowhead). RNA of four different regions of the membrane (A to D indicated on the right) were isolated from the membrane and size separated on a denaturing gel (6%) (right; nucleotide size marker on the left, bp). (**B**) Autoradiographs showing Rrm4-Gfp, Grp1-Gfp and Gfp in complex with RNA after UV-C irradiation at 0, 160, 320, 480 and 640 mJ/cm^2^. Corresponding Western blots using anti-Gfp are shown below. Arrowheads indicate the expected molecular weight of the proteins (Rrm4-Gfp, 112 kDa; Grp1-Gfp, 45 kDa; Gfp, 27 kDa). After each irradiation step, the cells were mixed. Note that increased UV-C irradiation in combination with slow processing due to long time intervals was particularly harmful for the Rrm4 protein, which was completely degraded after four minutes of UV-C irradiation. Putative degradation products are marked by asterisks. (**C**) Autoradiographs showing Rrm4-Gfp, Grp1-Gfp and Gfp in complex with RNA after single UV-C irradiation at 0, 100, 200, 300 or 400 mJ/cm^2^ This time, mixing breaks were omitted and cells were harvested as quickly as possible. Corresponding Western blots are shown below. Labelling as above. We chose 200 mJ/cm^2^ as optimal UV-C irradiation dose, since the amount of unspecific Gfp-RNA complexes increased at higher doses. Arrowheads and asterisks as in (B). (**D**) Amplification of the Rrm4, Grp1 and Gfp derived cDNA libraries with different numbers of PCR cycles (between 18 and 24; ctrl, control without template cDNA). The PCR products were separated on a native gel (6%) and stained with SYBR green I (nucleotide size marker on the left, bp). The size of the cDNA insert together with the adapters (cDNA insert = 20-30 nt; L3 adapter, RT-primer and P3/P5 Solexa primers = 128 nt) is expected to be ~ 150-160 nt after amplification.

**Figure EV3.**
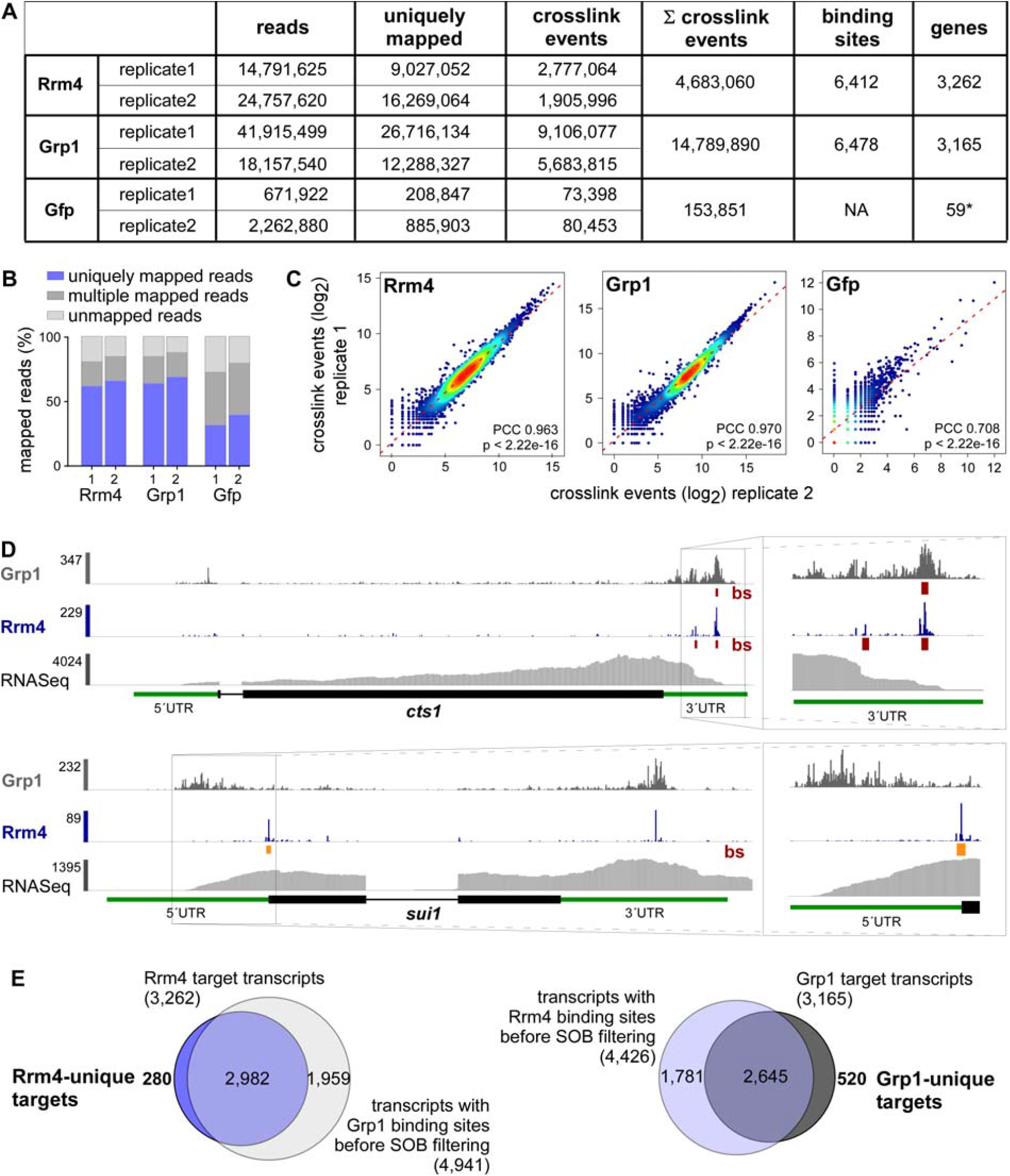
Comparative iCLIP procedure results in high-quality dataset. (**A**) Summary of the iCLIP libraries including initial number of sequencing reads, uniquely mapped reads, crosslink events (xlinks) for both replicates for Rrm4, Grp1 and Gfp. In addition, sum of crosslink events as well as resulting binding sites and target transcripts are given for merged replicates. Gfp binding sites were only filtered for reproducibility but not for relative signal intensity (SOB; see Material and methods). (**B**) Stacked bar chart showing percentage of reads mapping to a unique, multiple (multiple mapping) or no location (unmapped) in the *U. maydis* genome for Rrm4, Grp1 and Gfp. (**C**) Scatter plot comparing number of crosslink events per gene from two independent replicate experiments for Rrm4, Grp1 and Gfp (PCC, Pearson correlation coefficient). (**D**) Genome browser views of Rrm4 and Grp1 iCLIP events as well as RNASeq data of *cts1* (UMAG_10419) and *rrm4* (UMAG_10836). Visualisation as in Fig 3C. (**E**) Venn diagram to identify target transcripts that are uniquely bound by Rrm4 (left) or Grp1 (right). Unique target transcripts (numbers given in bold) are selected only if they show no evidence of binding by the other RBP (considering all binding sites before SOB filtering, see Materials and methods).

**Figure EV4.**
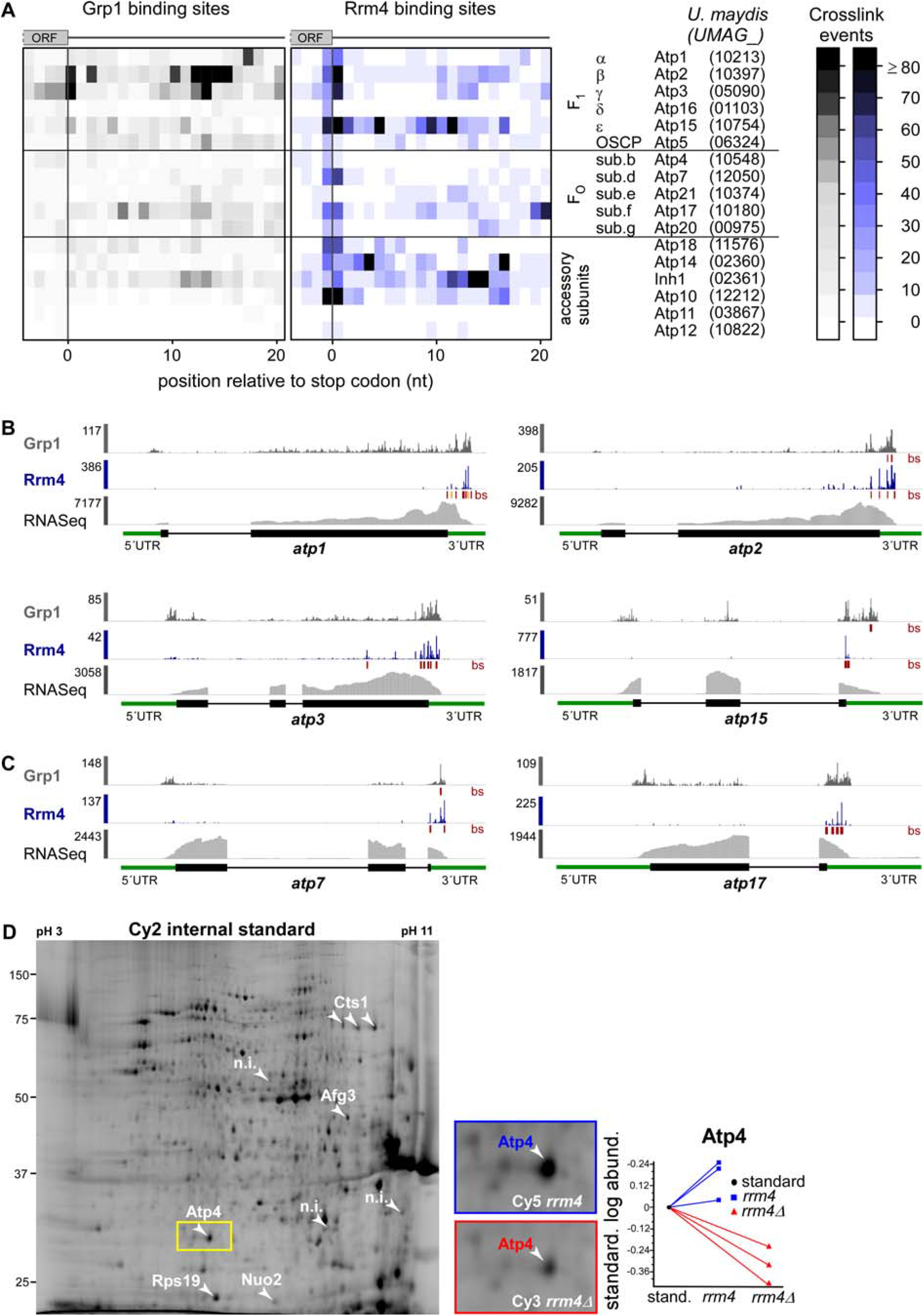
Accumulation of crosslink events at stop codons of mRNAs encoding subunits of the mitochondrial F_O_F_1_-ATPase. (**A**) Heatmap of crosslink events of Grp1 (left) and Rrm4 (right) in a window around the stop codons (position 0 = first position of 3’ UTR) of mRNAs encoding subunits of the mitochondrial F_O_F_1_-ATPase (nomenclature and gene identifiers for *U. maydis* on the right). Crosslink events per nucleotide are represented by a colour scale (right). (**B, C**) iCLIP data of Rrm4 and Grp1 as well as RNASeq data across selected mRNAs of the F1 subcomplex (**B**) and FO subcomplex (**C**) that carry an Rrm4 binding site precisely at the stop codon. Visualisation as in Fig 3C. (**D**) Cy2 image of a representative differential gel electrophoresis (DIGE) analysis comparing membrane-associated proteins of wild type (blue, Cy5-labelled) and *rrm4Δ* hyphae (red, Cy3-labelled; size marker on the left, pH range at the top; 6 h.p.i.). Protein variants exhibiting at least 2.5-fold differences in protein amounts are indicated by numbered arrowheads (taken from earlier study) [18]. The area marked with a yellow rectangle is enlarged on the right. The corresponding standardised logarithmic protein abundances for the Atp4 spot obtained from three biological replicates are given on the right (internal standard set to 0).

**Figure EV5.**
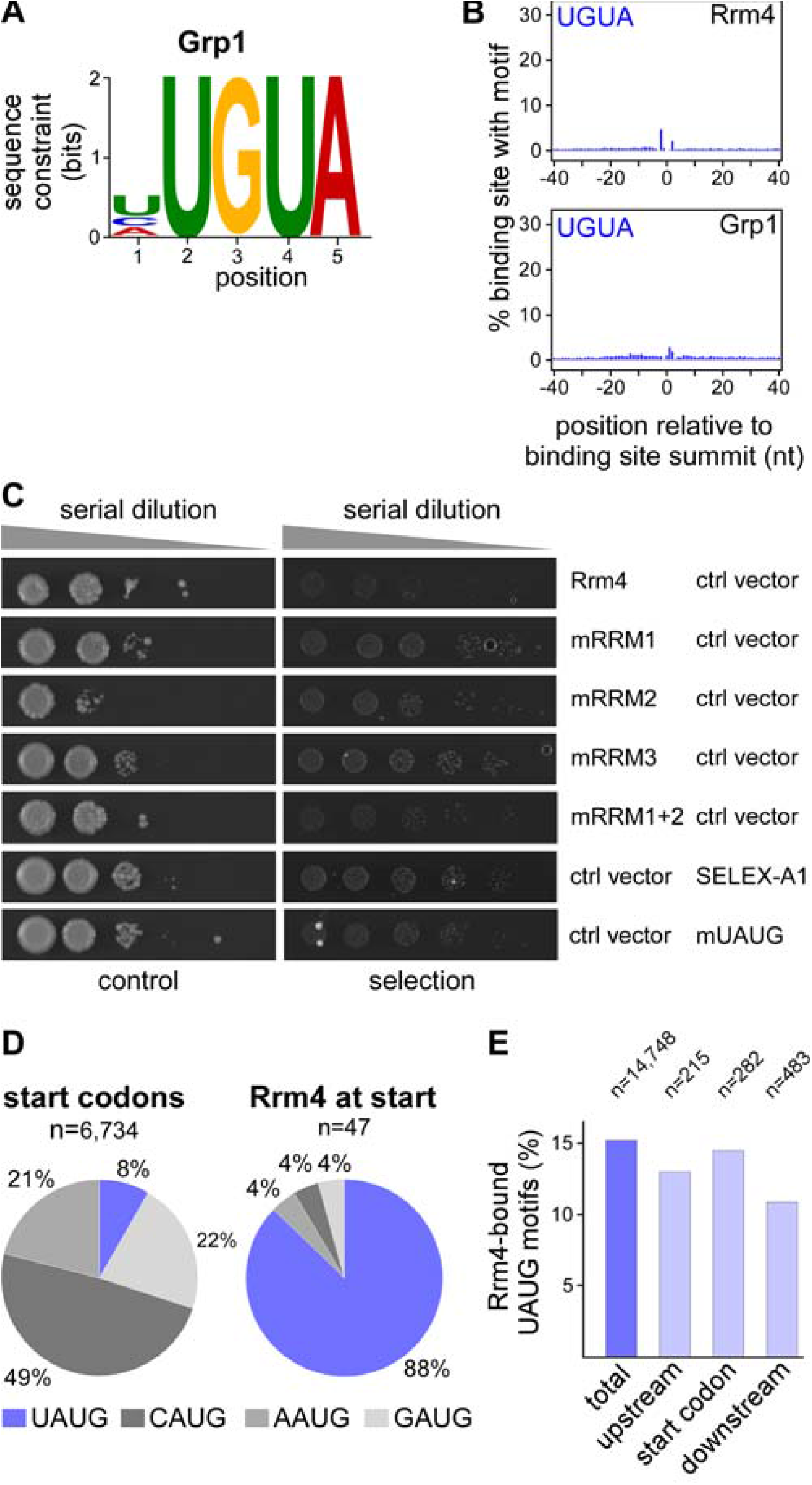
Control experiments for the yeast three-hybrid analysis. (**A**) Logo representation of the most enriched sequence motif at Grp1 binding sites. At each position, the height of the stack is proportional to the information content, while the relative height of each nucleotide within the stack represents its relative frequency at this position. (**B**) Frequency of UGUA around Rrm4 and Grp1 binding sites. Shown is the percentage of binding sites that harbour an UGUA starting at a given position in an 81-nt window around the binding site summit. Representation as in Fig 5B. (**C**) Colony growth on control and selection plates of yeast cells expressing protein and RNA hybrids indicated on the right. RNA binding is scored by growth on selection plates (SC –his +3-AT, 3-amino-1,2,4-triazole). This control experiment demonstrates that growth on selection plates (see Fig 5F) depends the presence of Rrm4 variant and cognate hybrid RNA. mRRMx, Rrm4 variants harbouring mutations in RRM 1, 2, 3 or 1 and 2. (**D**) Relative occurrence of NAUG sequence context for all (left) and Rrm4-bound (right) start codons. CAUG fits to the Kozak sequence in eukaryotes [81]. The fraction of start codons coinciding with the Rrm4 recognition motif UAUG is shown in blue. This sequence context was strongly enriched among the Rrm4-bound target mRNAs, whereas it comprises only 8% of all annotated start codons in the *U. maydis* genome. (**E**) UAUG motifs at start codons were not more frequently bound than UAUG motifs in the surrounding sequence (see Materials and methods). Out of a total of 14,748 UAUG motifs in expressed transcripts 15.2% are bound by Rrm4. Similarly, 14.5% of the 282 UAUG motifs directly at start codons are bound by Rrm4. This is only marginally more than at 215 and 483 UAUG motifs within 100 nt upstream and downstream of the start codons, out of which 13.0% and 11.0% are bound, respectively.

## References

1. Holt CE, Bullock SL (2009) Subcellular mRNA localization in animal cells and why it matters. Science 326: 1212–1216

2. Martin KC, Ephrussi A (2009) mRNA localization: gene expression in the spatial dimension. Cell 136: 719–730

3. Eliscovich C, Singer RH (2017) RNP transport in cell biology: the long and winding road. Current opinion in cell biology 45: 38–46

4. Mofatteh M, Bullock SL (2017) SnapShot: Subcellular mRNA localization. Cell 169: 178.e1

5. Baumann S, Pohlmann T, Jungbluth M, Brachmann A, Feldbrügge M (2012) Kinesin-3 and dynein mediate microtubule-dependent co-transport of mRNPs and endosomes. J Cell Sci 125: 2740–2752

6. Baumann S, König J, Koepke J, Feldbrügge M (2014) Endosomal transport of septin mRNA and protein indicates local translation on endosomes and is required for correct septin filamentation. EMBO Rep 15: 94–102

7. Egan MJ, McClintock MA, Reck-Peterson SL (2012) Microtubule-based transport in filamentous fungi. Curr Opin Microbiol 15: 637–645

8. Steinberg G (2014) Endocytosis and early endosome motility in filamentous fungi. Curr Opin Microbiol 20: 10–18

9. Guimaraes SC, Schuster M, Bielska E, Dagdas G, Kilaru S, Meadows BR, Schrader M, Steinberg G (2015) Peroxisomes, lipid droplets, and endoplasmic reticulum “hitchhike” on motile early endosomes. J Cell Biol 211: 945–954

10. Salogiannis J, Egan MJ, Reck-Peterson SL (2016) Peroxisomes move by hitchhiking on early endosomes using the novel linker protein PxdA. The Journal of cell biology 212: 289–296

11. Salogiannis J, Reck-Peterson SL (2016) Hitchhiking: a non-canonical mode of microtubule-based transport. Trends in cell biology 27: 141–150

12. Haag C, Pohlmann T, Feldbrugge M (2017) The ESCRT regulator Did2 maintains the balance between long-distance endosomal transport and endocytic trafficking. PLoS Genet 13: e1006734

13. Haag C, Steuten B, Feldbrügge M (2015) Membrane-coupled mRNA trafficking in fungi. Annu Rev Microbiol 69: 265–281

14. Vollmeister E, Feldbrügge M (2010) Posttranscriptional control of growth and development in *Ustilago maydis*. Curr Opin Microbio 13: 693–699

15. Becht P, Vollmeister E, Feldbrügge M (2005) Role for RNA-binding proteins implicated in pathogenic development of *Ustilago maydis*. Euk Cell 4: 121–133

16. Becht P, König J, Feldbrügge M (2006) The RNA-binding protein Rrm4 is essential for polarity in *Ustilago maydis* and shuttles along microtubules. J Cell Sci 119: 4964–4973

17. König J, Baumann S, Koepke J, Pohlmann T, Zarnack K, Feldbrügge M (2009) The fungal RNA-binding protein Rrm4 mediates long-distance transport of *ubi1* and *rho3* mRNAs. EMBO J 28: 1855–1866

18. Koepke J, Kaffarnik F, Haag C, Zarnack K, Luscombe NM, König J, Ule J, Kellner R, Begerow D, Feldbrügge M (2011) The RNA-binding protein Rrm4 is essential for efficient secretion of endochitinase Cts1. Mol Cell Proteom 10: M111.011213 1–15

19. Zander S, Baumann S, Weidtkamp-Peters S, Feldbrügge M (2016) Endosomal assembly and transport of heteromeric septin complexes promote septin cytoskeleton formation. J Cell Sci 129: 2778–2792

20. Zhu X, Buhrer C, Wellmann S (2016) Cold-inducible proteins CIRP and RBM3, a unique couple with activities far beyond the cold. Cellular and molecular life sciences: CMLS 73: 3839–3859

21. Kang H, Park SJ, Kwak KJ (2013) Plant RNA chaperones in stress response. Trends Plant Sci 18: 100–106

22. Brachmann A, Weinzierl G, Kämper J, Kahmann R (2001) Identification of genes in the bW/bE regulatory cascade in *Ustilago maydis*. Mol Microbiol 42: 1047–1063

23. Imai K, Noda Y, Adachi H, Yoda K (2005) A novel endoplasmic reticulum membrane protein Rcr1 regulates chitin deposition in the cell wall of Saccharomyces cerevisiae. J Biol Chem 280: 8275–84

24. Ram AF, Klis FM (2006) Identification of fungal cell wall mutants using susceptibility assays based on Calcofluor white and Congo red. Nat Protoc 1: 2253–2256

25. Baumann S, Takeshita N, Grün N, Fischer R, Feldbrügge M (2015) Live cell imaging of endosomal trafficking in fungi. In Methods in Mol Biol: Membrane trafficking, Tang BL (ed) pp 347–363. New York: Springer

26. Merzlyak EM, Goedhart J, Shcherbo D, Bulina ME, Shcheglov AS, Fradkov AF, Gaintzeva A, Lukyanov KA, Lukyanov S, Gadella TW, et al. (2007) Bright monomeric red fluorescent protein with an extended fluorescence lifetime. Nature methods 4: 555557

27. Campbell RE, Tour O, Palmer AE, Steinbach PA, Baird GS, Zacharias DA, Tsien RY (2002) A monomeric red fluorescent protein. Proc Natl Acad Sci U S A 99: 7877–7882

28. Hogan DJ, Riordan DP, Gerber AP, Herschlag D, Brown PO (2008) Diverse RNA-binding proteins interact with functionally related sets of RNAs, suggesting an extensive regulatory system. PLoS Biol 6: e255

29. König J, Zarnack K, Rot G, Curk T, Kayikci M, Zupan B, Turner DJ, Luscombe NM, Ule J (2010) iCLIP reveals the function of hnRNP particles in splicing at individual nucleotide resolution. Nat Struct Mol Biol 17: 909–915

30. Huppertz I, Attig J, D’Ambrogio A, Easton LE, Sibley CR, Sugimoto Y, Tajnik M, König J, Ule J (2014) iCLIP: protein-RNA interactions at nucleotide resolution. Methods 65: 274–287

31. Kämper J, Kahmann R, Bölker M, Ma LJ, Brefort T, Saville BJ, Banuett F, Kronstad JW, Gold SE, Müller O, et al. (2006) Insights from the genome of the biotrophic fungal plant pathogen *Ustilago maydis*. Nature 444: 97–101

32. Chabanon H, Mickleburgh I, Burtle B, Pedder C, Hesketh J (2005) An AU-rich stem-loop structure is a critical feature of the perinuclear localization signal of c-myc mRNA. Biochem J 392: 475–483

33. Levadoux M, Mahon C, Beattie JH, Wallace HM, Hesketh JE (1999) Nuclear import of metallothionein requires its mRNA to be associated with the perinuclear cytoskeleton. The Journal of biological chemistry 274: 34961–34966

34. Straube A, Enard W, Berner A, Wedlich-Söldner R, Kahmann R, Steinberg G (2001) A split motor domain in a cytoplasmic dynein. EMBO J 20: 5091–5100

35. Bailey TL (2011) DREME: motif discovery in transcription factor ChIP-seq data. Bioinformatics 27: 1653–1659

36. König J, Julius C, Baumann S, Homann M, Göringer HU, Feldbrügge M (2007) Combining SELEX and yeast three-hybrid system for *in vivo* selection and classification of RNA aptamers. RNA 13: 614–622

37. Lécuyer E, Yoshida H, Parthasarathy N, Alm C, Babak T, Cerovina T, Hughes TR, Tomancak P, Krause HM (2007) Global analysis of mRNA localization reveals a prominent role in organizing cellular architecture and function. Cell 131: 174–187

38. Jambor H, Surendranath V, Kalinka AT, Mejstrik P, Saalfeld S, Tomancak P (2015) Systematic imaging reveals features and changing localization of mRNAs in Drosophila development. eLife 4

39. Cajigas IJ, Tushev G, Will TJ, tom Dieck S, Fuerst N, Schuman EM (2012) The local transcriptome in the synaptic neuropil revealed by deep sequencing and high-resolution imaging. Neuron 74: 453–466

40. Rangaraju V, Tom Dieck S, Schuman EM (2017) Local translation in neuronal compartments: how local is local? EMBO Rep 18: 693–711

41. Calabretta S, Richard S (2015) Emerging roles of disordered sequences in RNA-binding proteins. Trends Biochem Sci 40: 662–672

42. Ying Y, Wang XJ, Vuong CK, Lin CH, Damianov A, Black DL (2017) Splicing activation by Rbfox requires self-aggregation through Its tyrosine-rich domain. Cell 170: 312-323 e10

43. Jung H, Yoon BC, Holt CE (2012) Axonal mRNA localization and local protein synthesis in nervous system assembly, maintenance and repair. Nat Rev Neurosci 13: 308–324

44. Kato M, Han TW, Xie S, Shi K, Du X, Wu LC, Mirzaei H, Goldsmith EJ, Longgood J, Pei J, et al. (2012) Cell-free formation of RNA granules: low complexity sequence domains form dynamic fibers within hydrogels. Cell 149: 753–767

45. Ciuzan O, Hancock J, Pamfil D, Wilson I, Ladomery M (2015) The evolutionarily conserved multifunctional glycine-rich RNA-binding proteins play key roles in development and stress adaptation. Physiol Plant 153: 1–11

46. Meyer K, Koster T, Nolte C, Weinholdt C, Lewinski M, Grosse I, Staiger D (2017) Adaptation of iCLIP to plants determines the binding landscape of the clock-regulated RNA-binding protein AtGRP7. Genome Biol 18: 204

47. Peretti D, Bastide A, Radford H, Verity N, Molloy C, Martin MG, Moreno JA, Steinert JR, Smith T, Dinsdale D, et al. (2015) RBM3 mediates structural plasticity and protective effects of cooling in neurodegeneration. Nature 518: 236–239

48. Bastide A, Peretti D, Knight JR, Grosso S, Spriggs RV, Pichon X, Sbarrato T, Roobol A, Roobol J, Vito D, et al. (2017) RTN3 is a novel cold-induced protein and mediates neuroprotective effects of RBM3. Curr Biol 27: 638–650

49. Kim JS, Park SJ, Kwak KJ, Kim YO, Kim JY, Song J, Jang B, Jung CH, Kang H (2007) Cold shock domain proteins and glycine-rich RNA-binding proteins from *Arabidopsis thaliana* can promote the cold adaptation process in *Escherichia coli*. Nucleic Acids Res 35: 506–516

50. Yano T, Lopez de Quinto S, Matsui Y, Shevchenko A, Shevchenko A, Ephrussi A (2004) Hrp48, a *Drosophila* hnRNPA/B homolog, binds and regulates translation of oskar mRNA. Dev Cell 6: 637–648

51. Huynh JR, Munro TP, Smith-Litiere K, Lepesant JA, St Johnston D (2004) The *Drosophila* hnRNPA/B homolog, Hrp48, is specifically required for a distinct step in osk mRNA localization. Dev Cell 6: 625–635

52. Goodrich JS, Clouse KN, Schupbach T (2004) Hrb27C, Sqd and Otu cooperatively regulate *gurken* RNA localization and mediate nurse cell chromosome dispersion in *Drosophila* oogenesis. Development 131: 1949–1958

53. König J, Zarnack K, Luscombe NM, Ule J (2012) Protein-RNA interactions: new genomic technologies and perspectives. Nat Rev Genet 13: 77–83

54. Van Nostrand EL, Pratt GA, Shishkin AA, Gelboin-Burkhart C, Fang MY, Sundararaman B, Blue SM, Nguyen TB, Surka C, Elkins K, et al. (2016) Robust transcriptome-wide discovery of RNA-binding protein binding sites with enhanced CLIP (eCLIP). Nature methods 13: 508–514

55. Zarnegar BJ, Flynn RA, Shen Y, Do BT, Chang HY, Khavari PA (2016) irCLIP platform for efficient characterization of protein-RNA interactions. Nature methods 13: 489–492

56. Doyle M, Kiebler MA (2011) Mechanisms of dendritic mRNA transport and its role in synaptic tagging. EMBO J 30: 3540–3552

57. Higuchi Y, Ashwin P, Roger Y, Steinberg G (2014) Early endosome motility spatially organizes polysome distribution. J Cell Biol 204: 343–357

58. Margeot A, Blugeon C, Sylvestre J, Vialette S, Jacq C, Corral-Debrinski M (2002) In *Saccharomyces cerevisiae*, *ATP2* mRNA sorting to the vicinity of mitochondria is essential for respiratory function. EMBO J 21: 6893–6904

59. Gehrke S, Wu Z, Klinkenberg M, Sun Y, Auburger G, Guo S, Lu B (2015) PINK1 and Parkin control localized translation of respiratory chain component mRNAs on mitochondria outer membrane. Cell Metab 21: 95–108

60. Gold VA, Chroscicki P, Bragoszewski P, Chacinska A (2017) Visualization of cytosolic ribosomes on the surface of mitochondria by electron cryo-tomography. EMBO Rep 18: 1786–1800

61. Lesnik C, Golani-Armon A, Arava Y (2015) Localized translation near the mitochondrial outer membrane: An update. RNA Biol 12: 801–809

62. Zhang Y, Chen Y, Gucek M, Xu H (2016) The mitochondrial outer membrane protein MDI promotes local protein synthesis and mtDNA replication. EMBO J 35: 1045–1057

63. Sen A, Cox RT (2016) Clueless is a conserved ribonucleoprotein that binds the ribosome at the mitochondrial outer membrane. Biol Open 5: 195–203

64. Pohlmann T, Baumann S, Haag C, Albrecht M, Feldbrügge M (2015) A FYVE zinc finger domain protein specifically links mRNA transport to endosome trafficking. eLife 4: 10.7554/eLife.06041

65. Halstead JM, Lionnet T, Wilbertz JH, Wippich F, Ephrussi A, Singer RH, Chao JA (2015) Translation. An RNA biosensor for imaging the first round of translation from single cells to living animals. Science 347: 1367–1671

66. Wu B, Eliscovich C, Yoon YJ, Singer RH (2016) Translation dynamics of single mRNAs in live cells and neurons. Science 352: 1430–1435

67. Graber TE, Hebert-Seropian S, Khoutorsky A, David A, Yewdell JW, Lacaille JC, Sossin WS (2013) Reactivation of stalled polyribosomes in synaptic plasticity. Proc Natl Acad Sci U S A 110: 16205–16210

68. Darnell JC, Van Driesche SJ, Zhang C, Hung KY, Mele A, Fraser CE, Stone EF, Chen C, Fak JJ, Chi SW, et al. (2011) FMRP stalls ribosomal translocation on mRNAs linked to synaptic function and autism. Cell 146: 247–261

69. Brachmann A, König J, Julius C, Feldbrügge M (2004) A reverse genetic approach for generating gene replacement mutants in *Ustilago maydis*. Mol Gen Genom 272: 216226

70. Shaner NC, Campbell RE, Steinbach PA, Giepmans BN, Palmer AE, Tsien RY (2004) Improved monomeric red, orange and yellow fluorescent proteins derived from *Discosoma* sp. red fluorescent protein. Nat Biotechnol 22: 1567–1572

71. Larkin MA, Blackshields G, Brown NP, Chenna R, McGettigan PA, McWilliam H, Valentin F, Wallace IM, Wilm A, Lopez R, et al. (2007) Clustal W and Clustal X version 2.0. Bioinformatics 23: 2947–2948

72. Baumann S, Zander S, Weidtkamp-Peters S, Feldbrügge M (2016) Live cell imaging of septin dynamics in Ustilago maydis. In Septins, Gladfelter AS (ed) pp 143–149. Elsevier Inc.

73. Rothbauer U, Zolghadr K, Muyldermans S, Schepers A, Cardoso MC, Leonhardt H (2008) A versatile nanotrap for biochemical and functional studies with fluorescent fusion proteins. Mol Cell Proteomics 7: 282–289

74. Dodt M, Roehr JT, Ahmed R, Dieterich C (2012) FLEXBAR-flexible barcode and adapter processing for next-generation sequencing platforms. Biology 1: 895–905

75. Dobin A, Davis CA, Schlesinger F, Drenkow J, Zaleski C, Jha S, Batut P, Chaisson M, Gingeras TR (2013) STAR: ultrafast universal RNA-seq aligner. Bioinformatics 29: 1521

76. Kucukural A, Ozadam H, Singh G, Moore MJ, Cenik C (2013) ASPeak: an abundance sensitive peak detection algorithm for RIP-Seq. Bioinformatics 29: 2485–2486

77. Sutandy FXR, Ebersberger S, Huang L, Busch A, Bach M, Kang HS, Fallmann J, Maticzka D, Backofen R, Stadler PF, et al. (2018) *In vitro* iCLIP-based modeling uncovers how the splicing factor U2AF2 relies on regulation by cofactors. Genome Res 28: 699–713

78. Ruepp A, Zollner A, Maier D, Albermann K, Hani J, Mokrejs M, Tetko I, Guldener U, Mannhaupt G, Münsterkötter M, et al. (2004) The FunCat, a functional annotation scheme for systematic classification of proteins from whole genomes. Nucleic Acids Res 32: 5539–5545

79. Vollmeister E, Haag C, Zarnack K, Baumann S, König J, Mannhaupt G, Feldbrügge M (2009) Tandem KH domains of Khd4 recognize AUACCC and are essential for regulation of morphology as well as pathogenicity in *Ustilago maydis*. RNA 15: 2206–2218

80. SenGupta DJ, Zhang B, Kraemer B, Pochart P, Fields S, Wickens M (1996) A three-hybrid system to detect RNA-protein interactions *in vivo*. Proc Natl Acad Sci U S A 93: 8496–8501

81. Kozak M (2005) Regulation of translation via mRNA structure in prokaryotes and eukaryotes. Gene 361: 13–37

