## Supplementary material for "The key protein of endosomal mRNP transport Rrm4 binds translational landmark sites of cargo mRNAs"

Appendix Table S4 ……………………………………………………………………..…05

**Appendix Table S1: *U. maydis* strains used in this study.**
eGfp, enhanced Gfp. UMa, internal reference number.

| Strain | Relevant genotype | Short description | Reference | UMa |
|---|---|---|---|---|
| AB33 | *a2 Pnar:bW2bE1* | expression of active b heterodimer under control of the $P_{nar1}$ promoter; strain grows filamentously upon changing the nitrogen source | Brachmann et al., 2001 | 133 |
| AB33rrm4Δ | *rrm4Δ* | carries a deletion of *rrm4* | Becht et al., 2006 | 273 |
| AB33rrm4G | *rrm4G* | expresses Rrm4 C-terminally fused to eGFP | Becht et al., 2006 | 274 |
| AB33pab1G | *pab1G* | expresses Pab1 C-terminally fused to eGfp | König et al., 2009 | 389 |
| AB33grp1Δ | *grp1Δ* | carries a deletion of *grp1* | This study | 380 |
| AB33grp1G | *grp1G* | expresses Grp1 C-terminally fused to eGfp | This study | 911 |
| AB33rrm4tR/grp1G | *grp1G/rrm4Δ* | co-expresses Rrm4 C-terminally fused to tagRfp and Grp1 C-terminally fused to eGfp | This study | 2169 |
| AB33pab1mC/grp1G | *pab1mC/grp1G* | co-expresses Pab1 C-terminally fused to mCherry and Grp1 C-terminally fused to eGfp | This study | 2200 |
| AB33pab1mC/rrm4G | *pab1mC/rrm4G* | co-expresses Pab1 C-terminally fused to mCherry and Rrm4 C-terminally fused to eGfp | König et al., 2009 | 554 |
| AB33rrm4G/grp1Δ | *rrm4G/grp1Δ* | expresses Rrm4 C-terminally fused to eGfp and carries a *grp1* deletion | This study | 880 |
| AB33grp1G/rrm4Δ | *grp1G/rrm4Δ* | expresses Grp1 C-terminally fused to eGfp and carries a *rrm4* deletion | This study | 1050 |
| AB33pab1G/rrm4Δ | *pab1G/rrm4Δ* | expresses Pab1 C-terminally fused to eGfp and carries a *rrm4* deletion | König et al., 2009 | 472 |
| AB33gfp | *gfp* | expresses *gfp* under control of the constitutive $P_{tef}$-promotor. The *gfp* construct is integrated ectopically into the *ipS*-locus. | Koepke et al., 2011 | 487 |
| AB33rrm4mR123tR/grp1G | *grp1G/rrm4Δ* | co-expresses Rrm4mR123 C-terminally fused to tagRfp and Grp1 C-terminally fused to eGfp | This study | 2012 |
| AB33rrm4mR123tR/pab1G | *pab1G/rrm4Δ* | co-expresses $Rrm4^{mR123}$ C-terminally fused to tagRfp and Pab1 C-terminally fused to eGfp | This study | 1200 |
| AB33 rrm4G/pab1mC/grp1Δ | *rrm4GT/pab1mC/grp1Δ* | co-expresses Rrm4 C-terminally fused to eGfp-tandem affinity purification tag and Pab1 C-terminally fused to mCherry, while carrying a *grp1* deletion | This study | 2156 |

**Appendix Table S2:** ***U. maydis* strains generated in this study.**
UMa and pUMa, internal reference numbers for strains and plasmids, respectively.

| Strain | UMa | Relevant genotype | Transformed plasmid | Locus | Progenitor strain |
|---|---|---|---|---|---|
| AB33grp1Δ | 380 | *grp1Δ* | pGrp1Δ-HygR (pUMa 848) | *grp1* | AB33 |
| AB33grp1G | 911 | *grp1G* | pGrp1G-NatR (pUMa 1595) | *grp1* | AB33 |
| AB33rrm4tR/grp1G | 2169 | *rrm4tR/grp1G* | pRrm4tR_G418R (pUMa 3004) | *rrm4* | AB33grp1G/rrm4Δ |
| AB33pab1mC/grp1G | 2200 | *pab1mC/grp1G* | pPab1mC_HygR (pUMa1208) | *pab1* | AB33grp1G |
| AB33rrm4G/grp1Δ | 880 | *rrm4G/grp1Δ* | pGrp1Δ-HygR (pUMa848) | *grp1* | AB33rrm4G |
| AB33grp1G/rrm4Δ | 1050 | *grp1G/rrm4Δ* | pRrm4Δ-HygR (pUMa1391) | *rrm4* | AB33grp1G |
| AB33rrm4mR123tR/grp1G | 2012 | *rrm4mR123tR/grp1G* | pRrm4mR123tR_G418R (pUMa2849) | *rrm4* | AB33grp1G/rrm4Δ |
| AB33rrm4mR123tR/pab1G | 1200 | *rrm4mR123tR/pab1G* | pRrm4mR123tR_G418R (pUMa2849) | *rrm4* | AB33pab1G |
| AB33 rrm4G/pab1mC/grp1Δ | 2156 | *rrm4G/grp1Δ/pab1mC* | pPab1mC-G418R (pUMa3052) | *pab1* | AB33 rrm4G/grp1Δ |

**Appendix Table S3: Plasmids generated in this study for *U. maydis*.**
pUMa, internal plasmid reference number.

| Plasmid | pUMa | Resistance cassette | Short description |
|---|---|---|---|
| pGrp1Δ-HygR | pUMa 848 | HygR (SfiI-insert of MF1hs) | Plasmid for generating deletion mutants of *grp1*. The hygromycin resistance cassette is flanked by the regions 0.9 kb upstream and 0.8 kb downstream of *grp1*. The flanking regions were amplified by PCR (UF: oSL217 ATGGTCTCGACTTGCGCG, oSL218 ttcggccatctaggccACGTAAACTTTGGCGGCC), (DF: oSL219, oSL220) using UM521 wild type DNA. |
| pGrp1G-NatR | pUMa 1595 | NatR (SfiI-insert of pMF5-1n) | Plasmid for generating eGfp-fusions of *grp1*. The eGfp cassette, containing the $T_{nos}$ terminator and the nourseothricin resistance cassette is flanked by a 2.3 kb upstream region, containing the entire *grp1* ORF, and a 0.8 kb downstream region. Both regions were amplified by PCR using genomic UM521 wild type DNA as template (UF: oSL865 GTCATTTAAATGGTCTCGACTTGCGCG, oSL863 CCATGGTGGCCGCGTTGGCCGCCTGGCTCTGTCCGTTGTATC; 2.3 kb); DF (oSL864 TGAGGCCTGAGTGGCCACGACTCTTGTCGGGGC, oSL866 GCTATTTAAATGTAGCGTTGCCACTCAGC; 0.8 kb). |
| pRrm4tR_G418R | pUMa 3004 | G418R (SfiI-insert of pMF1-g) | Plasmid for generating tagRfp-3x myc tag fusions of tagRfp of *rrm4*. The tagRfp cassette contains the $T_{nos}$ terminator and a G418 resistance cassette. It is flanked by a 3.2 kb upstream region, containing the entire 2.4 kb *rrm4* ORF, and the 2 kb immediately downstream of *rrm4*. The plasmid is a derivate of pRrm4G-NatR (Becht et al., 2006), with eGfp being exchanged for tagRfp. |
| pRrm4mR123tR_G418R | pUMa2849 | | Plasmid for generating tagRfp-3x myc tag fusions of *rrm4*$^{mR123}$. pRrm4tR_G418R, but carrying mutations in the RNP1 region of each RRM. |
| pPab1mK-G418R | pUMa3052 | | Plasmid for generating $mKate_2$-HA tag fusions of *pab1*. The mKate2 cassette contains the $T_{nos}$ terminator and a G418 resistance cassette. The cassette is flanked by a 3 kb upstream region, containing the 2 kb *pab1* ORF, and the 1 kb downstream of *pab1*. The plasmid is a derivate of pPab1G-NatR (König et al., 2009). |

**Appendix Table S4: Plasmids used for yeast three-hybrid system.**
pUMa, internal plasmid reference number.

| Plasmid | pUMa | Gene | Short description |
|---|---|---|---|
| pACTII-Ade | 399 | | Plasmid for the expression of hybrid proteins. Contains a Gal4 activation domain, to which sequences of interest can be fused. For positive selection this plasmid carries the *LEU2* and *ADE2* auxotrophy markers. Served as vector control |
| pRrm4-Gfp-pACTII-Ade | 427 | *rrm4* | Plasmid for expressing an Rrm4 hybrid protein, fused N-terminally to a Gal4 activation domain and C-terminally to eGfp. The plasmid is derived from pACTII-Ade (König et al., 2007). |
| pIII MS2-5 | 494 | | Plasmid for expressing RNA-hybrids. Contains a *RPR1* leader, followed by two MS2 coat protein binding sites, a junk sequence flanked by a ClaI & AscI restriction sites, and a *RPR1* terminator. For positive selection this plasmid carries the *URA3* auxotrophy marker. Sequences can be cloned via the ClaI/ AscI restriction sites. Served as vector control. |
| pRrm4-mR3-Gfp-pACTII-Ade | 526 | *rrm4* | Plasmid for expressing the $Rrm4^{mR3}$ hybrid protein. Similar to pRrm4-gfp-pACTII-Ade, but carries a mutation in the RNP1 region of RRM3 (König et al., 2007) |
| pIII-MS2-5-Sel13 | 550 | | Plasmid for expressing the RNA-hybrid containing the SELEX aptamer A1. SELEX library sequences were inserted into pIII MS2-4 via ClaI/AscI (König et al., 2007). |
| pRrm4-mR1-Gfp-pACTII-Ade | 581 | *rrm4* | Plasmid for expressing the $Rrm4^{mR1}$ hybrid protein. Similar to pRrm4-gfp-pACTII-Ade, but carries a mutation in the RNP1 region of RRM1 (König et al., 2007). |
| pRrm4-mR12-Gfp-pACTII-Ade | 591 | *rrm4* | Plasmid for expressing the $Rrm4^{mR12}$ hybrid protein. Similar to pRrm4-gfp-pACTII-Ade, but carries mutations in the RNP1 regions of RRM1 and RRM2. |
| pRrm4-mR2-Gfp-pACTII-Ade | 594 | *rrm4* | Plasmid for expressing the $Rrm4^{mR2}$ hybrid protein. Similar to pRrm4-gfp-pACTII-Ade, but carries a mutation in the RNP1 region of RRM2 (König et al., 2007) |
| pIII MS2-5 A1- mUAUG | 1937 | | Plasmid for expressing the RNA-hybrid containing SELEX aptamer $A1^{mUAUG}$. TATGC was mutated to TCTCA by inverse PCR using oligos oRL750 GTGTGAAAGATCCGAAGTTTCTCACGGCACGCGCTGGCGCTG & oRL751 CAGCGCCAGCGCGTGCCGTGAGAAACTTCGGATCTTTCACAC and pIII-MS2-5-Sel13 as template. |
